# Characterization of new thermophilic antibiotic resistance markers

**DOI:** 10.64898/2025.12.15.694393

**Authors:** Fernanda Souza Lopes, Renato Vicentini, Edson Kim, Nandhini Ashok, Adam M. Guss, Lee R. Lynd, Luana Walravens Bergamo, Daniel G. Olson

## Abstract

The genetic engineering of thermophilic bacteria is constrained by limited availability of thermostable antibiotic resistance markers for selection. *Clostridium thermocellum*, a promising candidate for consolidated bioprocessing of lignocellulosic biomass, requires reliable selection systems that function at elevated temperatures. Here, we systematically evaluated antibiotic susceptibility profiles and identified novel resistance markers for this thermophile through bioinformatic screening and experimental validation. We screened 823 thermophilic genomes against the Comprehensive Antibiotic Resistance Database, identifying 1,115 antibiotic resistance genes. From these, we selected candidates with highest homology to resistance determinants for rifampicin, tetracycline, erythromycin, thiamphenicol, and neomycin. We identified three novel antibiotic resistance systems that function in this organism: tetracycline/*tet(45)*, erythromycin/*cmeC*, and rifampicin/*rbpA*. Of these, the rifampicin/rbpA provided the highest selection range , > 10,000-fold. Our results establish *rbpA* as an outstanding selectable marker for thermophilic genetic engineering and provide a validated workflow for discovering thermostable resistance determinants in high-temperature microorganisms.

**Importance:** Thermophilic bacteria like *Clostridium thermocellum* hold tremendous potential for sustainable biofuel production from plant biomass, but their genetic manipulation has been severely limited by the lack of selection markers that work at high temperatures. Many existing antibiotic resistance systems do not function at thermophilic temperatures, and many approaches to genetic manipulation require multiple antibiotic resistance markers. Currently only two markers are available for *C. thermocellum*, and only one (*cat*) functions well. The newly-developed *rbpA* marker functions well in *C. thermocellum* and is likely to provide dramatic new opportunities for engineering thermophilic host organisms.

## 1. Introduction

*Clostridium thermocellum* (also referred to as *Acetivibrio thermocellus*) is an anaerobic, thermophilic bacterium that has attracted considerable interest for the consolidated bioprocessing (CBP) of lignocellulosic biomass. Its native capacity to both hydrolyze cellulose and ferment the resulting sugars into biofuels and other commodities makes it a promising candidate for cellulosic biofuel production (Lynd et al., 2002). Cellulose and hemicellulose represent the two most abundant organic materials on Earth, and their conversion into fuels and chemicals has strong sustainability benefits, including potential carbon-neutral or even carbon-negative process pathways (Canadell and Schulze, 2014; Field et al., 2020).

Beyond its metabolic capabilities, *C. thermocellum* stands out for exceptionally rapid solubilization and growth on cellulosic substrates relative to many other cellulolytic microorganisms, which has established it as a leading model organism for CBP research (Paye et al., 2016; Xu et al., 2016). This thermophilic microorganism grows optimally at elevated temperatures (≈55–60 °C) and, when cultivated on cellulosic biomass, produces hydrogen, ethanol, and acetate among other fermentation products (Carere et al., 2008). Such physiological traits both enable faster biomass turnover and pose unique opportunities and challenges for industrial application and genetic manipulation. Developing efficient genetic tools for *C. thermocellum* thus depends not only on understanding its physiology but also on overcoming molecular constraints associated with selection systems at high temperatures. Despite an extensive body of work aimed at decoding its metabolism and developing genetic tools, there remains a pressing need for reliable selection markers (particularly antibiotic resistance markers) that function robustly under thermophilic conditions to support strain engineering and industrial strain development.

Antibiotics are small bioactive molecules naturally synthesized by microorganisms such as bacteria and fungi as part of their secondary metabolism, serving ecological roles in microbial competition and communication (Okada and Seyedsayamdost, 2017). These compounds exhibit diverse mechanisms of action, targeting essential cellular processes such as protein, RNA, and DNA synthesis. However, the genetic engineering of *C. thermocellum* and other thermophilic microorganisms has been constrained by the limited availability of thermostable antibiotics and their corresponding resistance markers. This limitation arises from the fact that most antibiotic-producing microorganisms are mesophilic bacteria and fungi, and many commonly used antibiotics lose activity at elevated temperatures (Noll and Vargas, 1997; Peteranderl et al., 1990). For example, ampicillin has a half-life of only 3.3 hours at 72 °C, making it unsuitable for stable selection in thermophilic hosts (Peteranderl et al., 1990). Nevertheless, certain antibiotics such as kanamycin, neomycin, chloramphenicol, and erythromycin have been reported to display sufficient stability under elevated temperatures, with kanamycin being the most thermostable among them (Peteranderl et al., 1990).

To explore the applicability of antibiotics in thermophilic systems, several studies have investigated antibiotic susceptibility in microorganisms adapted to high-temperature environments. For instance, Kortam et al. (2023) examined thermophilic bacteria isolated from hot spring waters and evaluated the susceptibility of *Bacillus licheniformis* isolate 113 and *Brevibacillus borstelensis* isolate 10 to 13 commercial antibiotics, including gentamicin, neomycin, tetracycline, rifamycin, streptomycin, and erythromycin (several of which overlap with the antibiotics tested in the present study). Their results demonstrated variable sensitivity among these thermophiles, indicating that despite the thermal instability of many antibiotics, some retain sufficient activity under elevated temperatures to serve as potential selection agents. Similarly, *Rhodothermus marinus*, a halophilic extreme thermophile, has been reported to be naturally sensitive to several 50S ribosomal subunit-targeting antibiotics such as erythromycin, chloramphenicol, and thiostrepton, as well as to rifampicin, which targets RNA polymerase (Silvia et al., 2021). This organism also yielded spontaneous resistant mutants to rifampicin and macrolides, suggesting that these antibiotics and their associated resistance determinants may be adapted for use as genetic markers in thermophilic bacteria. Together, these findings support the rationale for selecting erythromycin, gentamicin, streptomycin, spectinomycin, tetracycline, and rifampicin for assessment in *C. thermocellum*, as these antibiotics have demonstrated partial stability and biological activity in other thermophilic systems, making them promising candidates for thermostable selection marker development.

Antibiotic resistance genes are the most widely applied selection markers in microbial recombinant DNA technology. However, the lack of robust thermostable resistance determinants is considered a critical bottleneck for the genetic manipulation of thermophiles (Taylor et al., 2011; Zeldes et al., 2015). Shuttle vectors developed for *Escherichia coli* and *Bacillus* commonly incorporate resistance genes from mesophilic organisms, such as chloramphenicol and kanamycin resistance determinants originating from *Staphylococcus aureus* (Haima et al., 1987). Although progress has been made such as the successful introduction of a tetracycline resistance marker into a moderately thermophilic strain, and the demonstration that spectinomycin resistance functions in two thermophilic species, these systems often require low antibiotic concentrations, which increases the risk of spontaneous emergence of resistant colonies (Suzuki et al., 2018; Zhou et al., 2016).

Efforts to overcome these limitations include the directed evolution of resistance proteins from mesophilic origins, resulting in thermostable variants for antibiotics such as bleomycin, hygromycin, and thiostrepton (Brouns et al., 2005; Wada et al., 2016). Although such strategies highlight the potential of adapting mesophilic resistance mechanisms for application in thermophilic systems, the repertoire of validated thermostable selection markers remains small, necessitating further exploration.

Despite these advances, few systematic studies have examined minimum inhibitory concentration (MIC) values for thermophiles at elevated temperatures. Addressing this knowledge gap, the present study aimed to identify and evaluate novel antibiotic resistance markers that remain functional at elevated temperatures, with a particular focus on *C. thermocellum*. Specifically, we investigated the susceptibility of this bacterium to a diverse set of antibiotics with distinct mechanisms of action, assessed how temperature influences resistance profiles, and explored the applicability of these markers for future genetic engineering efforts. To achieve this, MIC assays were employed as the benchmark methodology, enabling a systematic comparison of antibiotic performance under thermophilic growth conditions, for both wild-type and mutant strains (i.e., with a resistant gene). The antibiotics tested included thiamphenicol, erythromycin, gentamicin, neomycin, streptomycin, spectinomycin, tetracycline, and rifampicin. Together, these efforts provide insights into the feasibility of developing thermostable selection systems and lay the groundwork for optimizing resistance markers in thermophiles for biotechnological applications.

## 2. Material and Methods

### 2.1 Antibiotic Sensitivity Assay and MIC Determination

The Minimum Inhibitory Concentration (MIC) for each antibiotic was determined by testing a range of concentrations to identify the lowest concentration that inhibited visible bacterial growth. All assays were performed using *Clostridium thermocellum* wild-type strain cultivated under strictly anaerobic conditions. The methodology was adapted from Wiegand et al. (2008) with modifications suitable for thermophilic anaerobes.

Bacterial stock cultures were grown in 10 mL of CTFUD liquid medium without antibiotics until reaching the mid-log phase. Then, serial dilutions (1:10, 1:100, and 1:1000) were prepared in pre-reduced CTFUD medium inside the anaerobic chamber. The final 1:1000 dilution was used to inoculate CTFUD agar plates (0.8% agar) supplemented with a wide range of concentrations for each antibiotic. Because the MICs were not known a priori, preliminary assays were designed to span a broad inhibitory range for each compound, extending below and above the expected MIC. The full set of concentrations tested for wild-type and plasmid-bearing strains is listed in Table 1. The composition of the CTFUD medium used in this study is shown in Table 2.

**Table 1.**
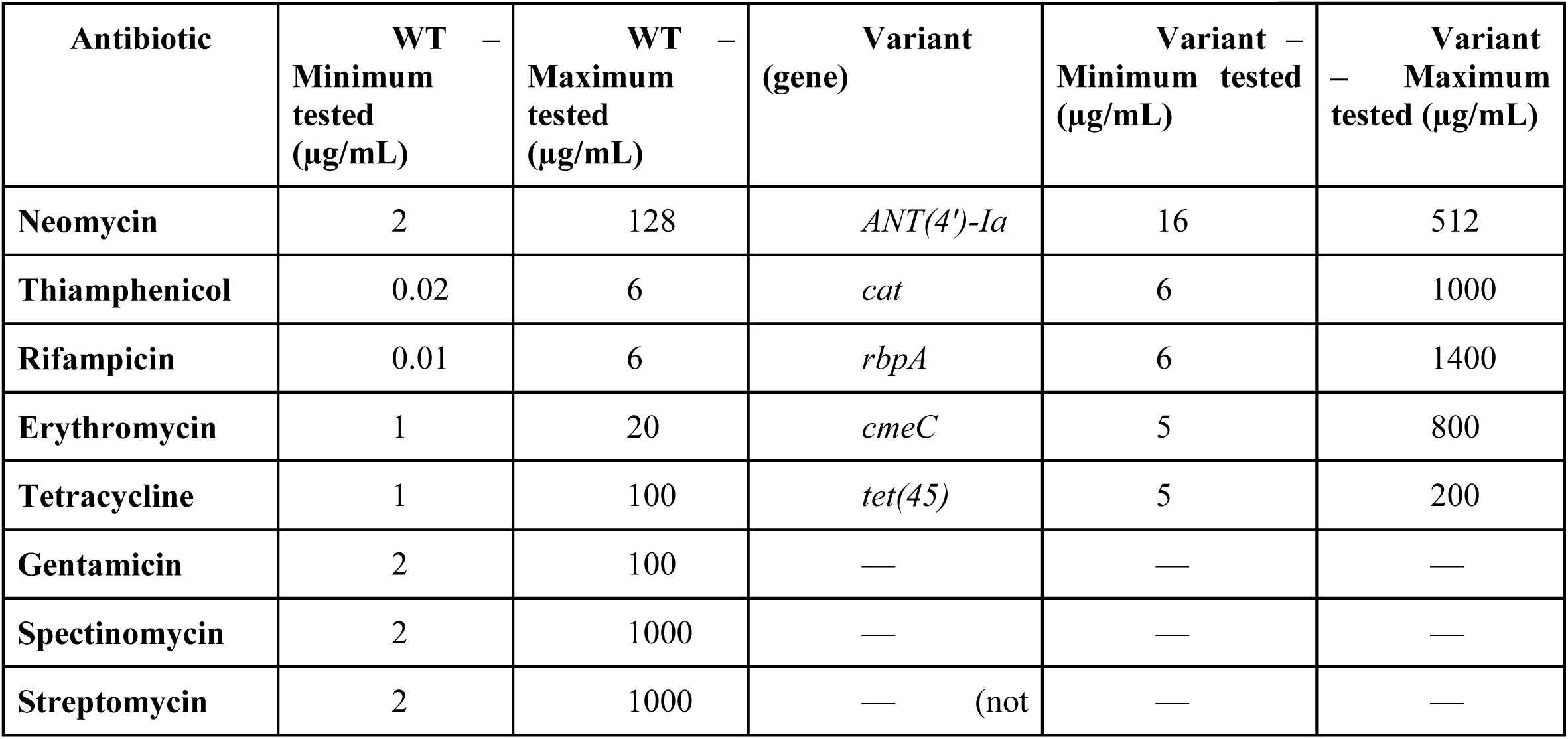
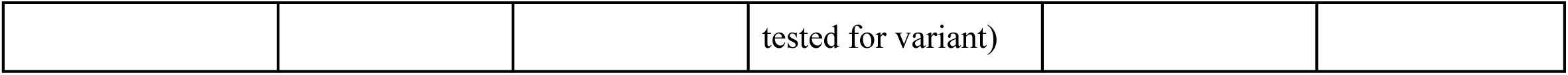
Antibiotic concentrations tested in MIC assays for wild-type *C. thermocellum* and plasmid-bearing strains. Values represent the lowest and highest concentrations included in the assay and do not correspond to MIC values.

**Table 2.**
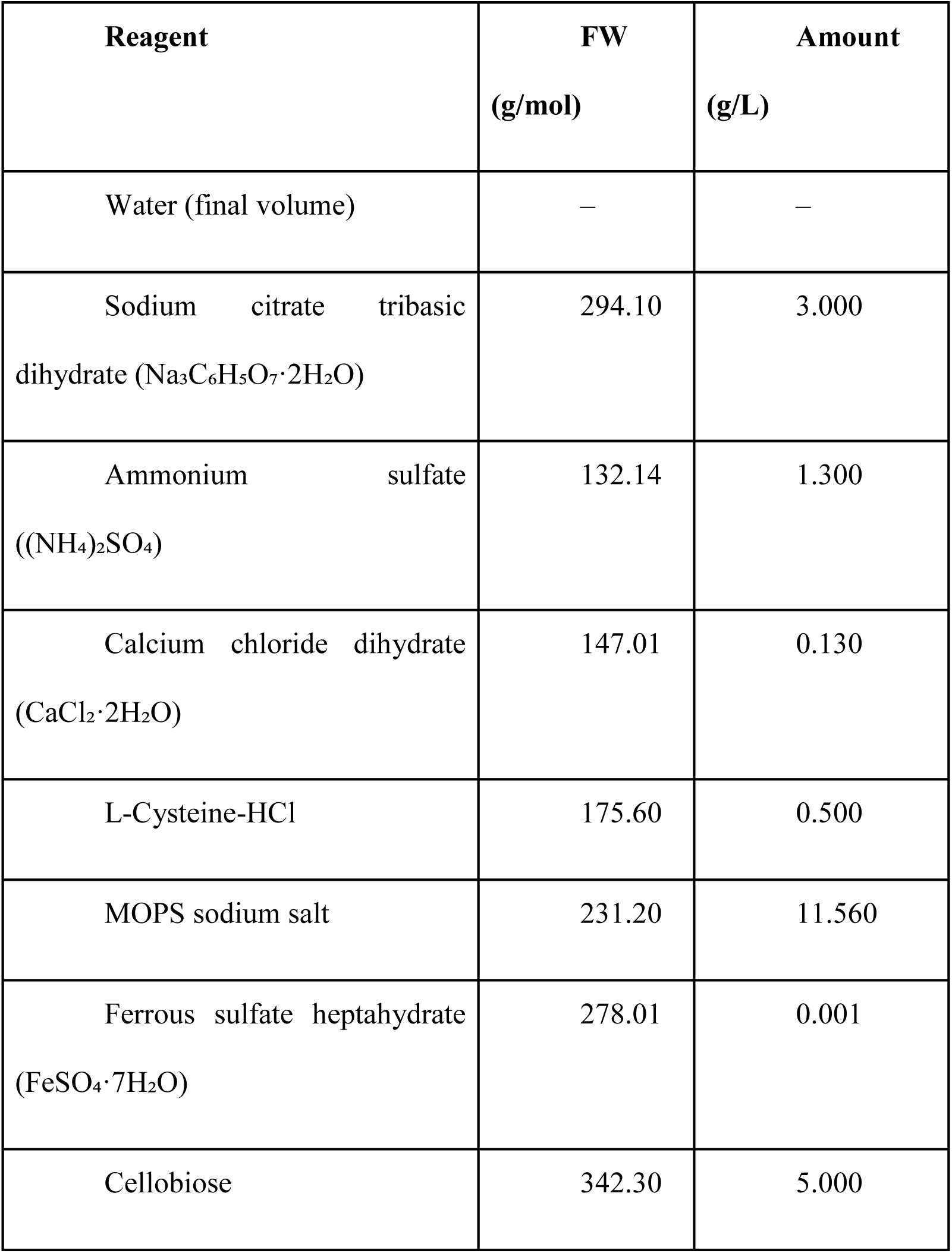
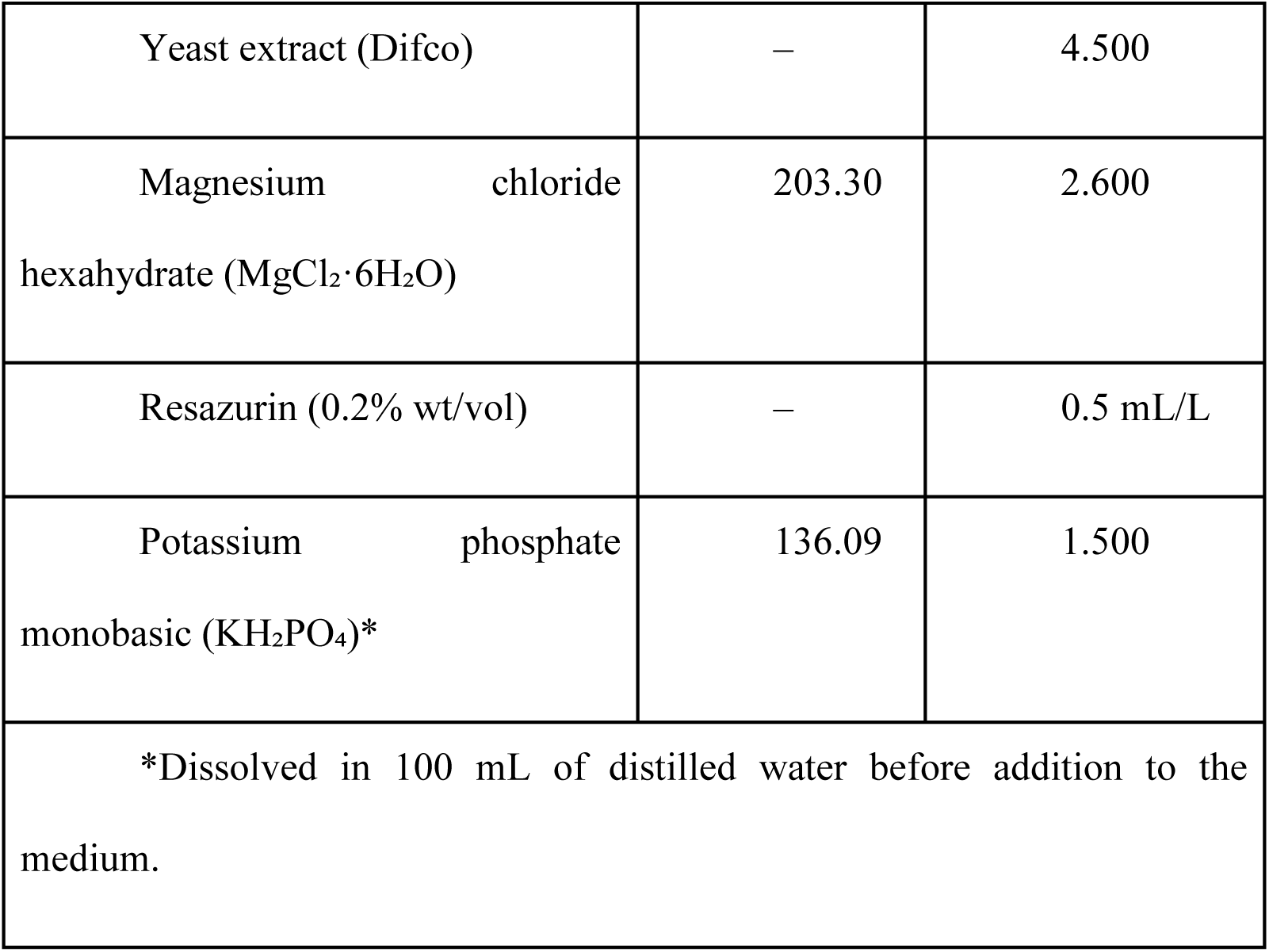
Composition of CTFUD medium used for growth and MIC assays.

For each concentration, duplicate plates were prepared: one set was incubated at 48°C and the other at 60°C to evaluate the effect of temperature on antibiotic susceptibility. Plates were incubated in anaerobic jars with AnaeroGen™ gas packs (Oxoid) to maintain an oxygen-free atmosphere and placed in a HERA THERM Thermo Scientific incubator at the respective temperatures.

A control plate without any antibiotic was included to confirm the viability of the bacterial culture and the adequacy of the anaerobic conditions. Thus, if bacterial growth was observed on the control plate but not on the antibiotic-containing plates, the inhibition was attributed to the antibiotic effect rather than a failure in the inoculation or growth conditions.

Each MIC determination was performed in triplicate, and each antibiotic concentration was tested in three replicate sectors on tripartite plates for each temperature.

Bacterial growth was visually assessed by colony formation on solid medium for 10 days, longer than a typical selection experiment that these markers will be used for (2-5 days). Evaluations were also recorded at 2 and 5 days post-inoculation (DPI) to monitor growth dynamics over time and to assess whether partial inhibition or delayed growth occurred under the different temperature conditions.

### 2.2 Plasmid Construction and Assembly Strategy

The plasmid pDGO143 (4135 bp) (Hon et al., 2016) was used as the initial template for genetic engineering (Figure 1). This vector served as the donor of the thiamphenicol antibiotic resistance gene(*cat*) and of the plasmid backbone used for downstream cloning. Plasmid DNA was digested with the restriction enzymes BglI + NsiI to excise the antibiotic resistance gene (*cat*) and linearize the backbone. This digestion excised non-target regions and generated a linearized template enriched for the *cat* locus, minimizing the likelihood of nonspecific amplification during PCR. In a parallel step, BamHI + HindIII were used to cleave the plasmid backbone, ensuring fragmentation of vector sequences and reducing the probability of unwanted background during Gibson Assembly. Restriction digestions were incubated at 37 °C for 2 h. Enzyme inactivation was performed according to manufacturer specifications: HindIII and NsiI at 80 °C for 20 min, BglI at 65 °C for 20 min. At the same time, the BamHI digest underwent silica-column purification to eliminate residual nuclease activity before PCR amplification.

**Figure 1.**
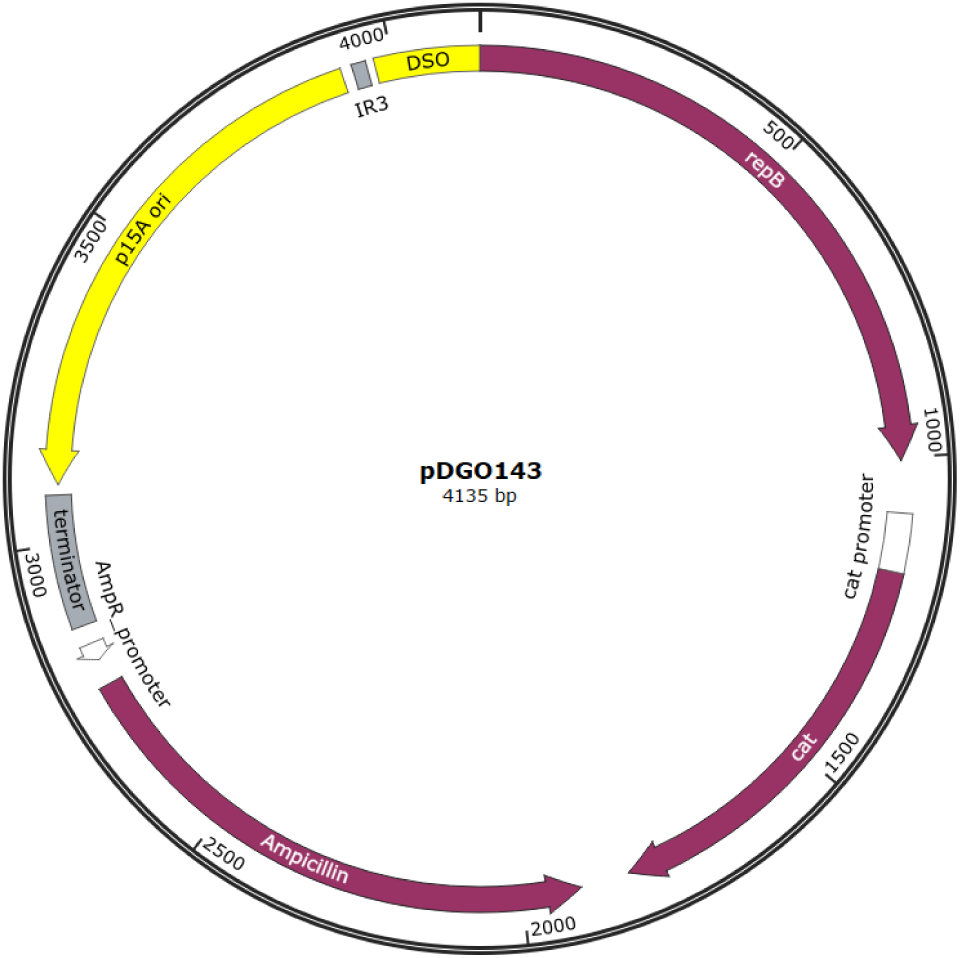
Structural map of plasmid pDGO143 (4,135 bp). The plasmid backbone carries the *E. coli* P15A medium-copy origin of replication (P15A ori) and the *C. thermocellum* pNW33N (Olson & Lee, 2012) origin of replication (double-strand origin (DSO) + *repB*). The *cat* gene conferring thiamphenicol resistance is expressed from the CAT promoter and flanked by promoter insulator elements and regulatory sequences to prevent transcriptional interference. The *AmpR* cassette, driven by the AmpR promoter, enables selection in *E. coli*.

Purification of the BamHI digestion was performed using a Cellco silica-based spin column (DPK-106L, Cellco Biotech do Brasil Ltda). The digested DNA was mixed with binding buffer, loaded onto the column, centrifuged to allow DNA adsorption to the silica membrane, washed to remove residual enzyme and salts, and finally eluted in nuclease-free water for downstream PCR.

Following restriction digestion, two independent high-fidelity PCR reactions were performed using Phusion™ High-Fidelity PCR Master Mix with HF Buffer (M0531S, New England Biolabs) to amplify (i) the *cat* gene and (ii) the pDGO143 backbone. For both reactions, the initial denaturation step was carried out at 98 °C for 2 min, followed by 35 amplification cycles. Each cycle consisted of denaturation at 98 °C for 10 s, annealing at 60°C, and extension at 75 °C. For the *cat* fragment, the annealing stage lasted 40 s and the extension step 45 s, whereas for the pDGO143 backbone, the annealing stage remained the same but the extension was prolonged to 1 min 55 s. A final extension step at 72 °C for 5 min was performed for both reactions to ensure complete amplification. All primers used for these amplifications and for subsequent plasmid construction steps are listed in Table 2, together with functional descriptions and notes on engineered sequence regions.

The promoter *pforA* was amplified from genomic DNA of *Thermoanaerobacterium saccharolyticum* LL1025 (Mai et al., 2006). This promoter was chosen because *pforA* is natively highly active in thermophiles and has been successfully used to drive expression of heterologous genes in *C. thermocellum*, improving pathway flux and product titers (Hon et al., 2018; Tian et al., 2019). The use of a promoter originating from a related thermophilic organism increases the likelihood of proper recognition by the host sigma factors and of correct transcriptional initiation at elevated temperatures, compared with mesophilic promoters. Amplification of *pforA* was performed using Phusion™ polymerase under the following cycling conditions: 98 °C for 2 min; 35 cycles of 98 °C for 10 s, 60 °C for 30 s, and 72 °C for 40 s; followed by a final extension at 72 °C for 5 min.

PCR products were purified using the Cellco PCR Purification Kit (DPK-106L, Cellco Biotech do Brasil Ltda) and quantified to determine pmol/µL using the formula:

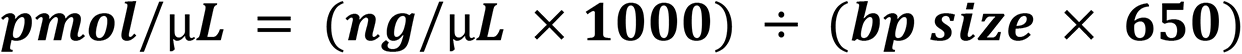

For Gibson Assembly, a molar ratio of 1:2:2 (vector: promoter: *cat*) was adopted. Based on the calculated pmol/µL values obtained from DNA quantification, the final assembly mixture contained 6.25 µL of backbone (0.1 pmol), 1.2 µL of the *pforA* promoter, 2 µL of the *cat* gene, 10 µL of Gibson Assembly® Master Mix (E2611L, New England Biolabs), and 0.55 µL of nuclease-free water, resulting in a final reaction volume of 20 µL. The mixture was incubated at 50 °C for 1 h and subsequently used for transformation into electrocompetent *E. coli* DH10β. For each electroporation, 1 µL of the Gibson reaction was added to the competent cells. A positive control (pUC) and a negative control (no-DNA) were included, as well as a pre-assembly background control in which 1 µL of the Gibson mix was transformed prior to the 50 °C incubation step to assess vector background. Following electroporation, 1 mL of SOC medium (B9020S,New England Biolabs) was added and cultures were incubated at 37 °C for 1 h with shaking before plating on LB agar supplemented with 100 ug/ml ampicillin (20 mL LB + 20 µL of 100 mg/mL ampicillin). Plates were incubated overnight at 37 °C.

Putative transformants were screened by colony PCR using primers PforA_fwd and cat_CDS_rev, and colonies yielding the expected amplicon size were inoculated into LB + ampicillin and grown overnight at 37 °C. Plasmid extraction was performed using the Fast-n-Easy Plasmid Miniprep Kit (DPK-104L, Cellco Biotech do Brasil Ltda), and the recovered plasmids were validated by EcoRV restriction digestion (2 h at 37 °C) followed by agarose gel electrophoresis.

To expand the applicability of the vector for selection in *C. thermocellum*, a second plasmid was generated in which the *cat* gene was replaced by the neomycin resistance determinant (*neo*), amplified from plasmid pDGO229 (also known as pLL1443 in the Lynd Lab plasmid collection), using the neo marker from plasmid pMU1592 (Olson, 2011). For this construct, the backbone was PCR-amplified from the validated pDGO143-PforA::*cat* plasmid, excluding the *cat* region, while *neo* was amplified separately. High-fidelity PCRs were performed with Phusion™ High-Fidelity PCR Master Mix (M0531L, New England Biolabs) using the following cycling conditions: for *neo*, 98 °C for 2 min followed by 35 cycles of 98 °C for 10 s, annealing at 60 °C for 40 s, and extension at 75 °C for 1 min, with a final extension step at 72 °C for 5 min; for the backbone lacking *cat*, 98 °C for 2 min followed by 35 cycles of 98 °C for 10 s, annealing at 60 °C for 45 s, and extension at 75 °C for 2 min, with a final extension of 72 °C for 5 min.

After quantification, the Gibson reaction was assembled using 2.5 µL of backbone (0.1 pmol), 0.74 µL of *neo* (0.2 pmol), 10 µL Gibson Assembly® Master Mix (E2611L, New England Biolabs), and 6.76 µL nuclease-free water in a final volume of 20 µL. The reaction was incubated at 50 °C for 1 h and transformed into *E. coli* DH10β following the same workflow used for the *cat* construct. Confirmed plasmids were subsequently used for the transformation of *C. thermocellum* and downstream MIC assays.

### 2.3 Bioinformatic Analysis

Thermophilic bacterial genomes were screened to identify potential antibiotic resistance determinants. A total of 823 thermophilic genomes were retrieved from the JGI Integrated Microbial Genomes & Microbiomes (IMG/M) database (https://img.jgi.doe.gov; Chen et al., 2023), using the advanced search filter *Phenotype Metadata → Temperature Range [thermophile* OR *hyperthermophile]*.

The corresponding genome assemblies were downloaded from the NCBI Genome database via command-line interface (CLI) tools and from the JGI Data Portal for bulk retrieval.

All genomes were analyzed against the Comprehensive Antibiotic Resistance Database (CARD), version 3.2.9 (Alcock et al., 2023), using the Resistance Gene Identifier (RGI) tool, version 5.2.0. Analyses were performed with the parameters -a DIAMOND --exclude_nudge --local, which applies the DIAMOND alignment algorithm (Buchfink et al., 2015) for high-throughput sequence comparison while excluding low-confidence or predicted hits.

To ensure biological relevance, downstream analyses focused only on the antibiotics previously tested in *C. thermocellum* sensitivity assays. Homology identity scores were used to select candidate genes with the highest similarity to CARD reference sequences for rifamycin, aminoglycoside, tetracycline, phenicol and macrolide drug classes (erythromycin, gentamicin, neomycin, streptomycin, spectinomycin, tetracycline, thiamphenicol and rifampicin). These resistance determinants were prioritized based on both their predicted thermostability and their relevance to the previously obtained antibiotic susceptibility profile of wild-type *C. thermocellum*. The corresponding coding sequences were synthesized and cloned into plasmid backbones to generate antibiotic-selectable vectors for transformation assays.

Table 4 provides an overview of all plasmids generated in this study, including Gibson-assembled constructs derived from pDGO143 and pDGO229, as well as plasmids carrying synthesized resistance genes selected through bioinformatic screening.

**Table 3.**
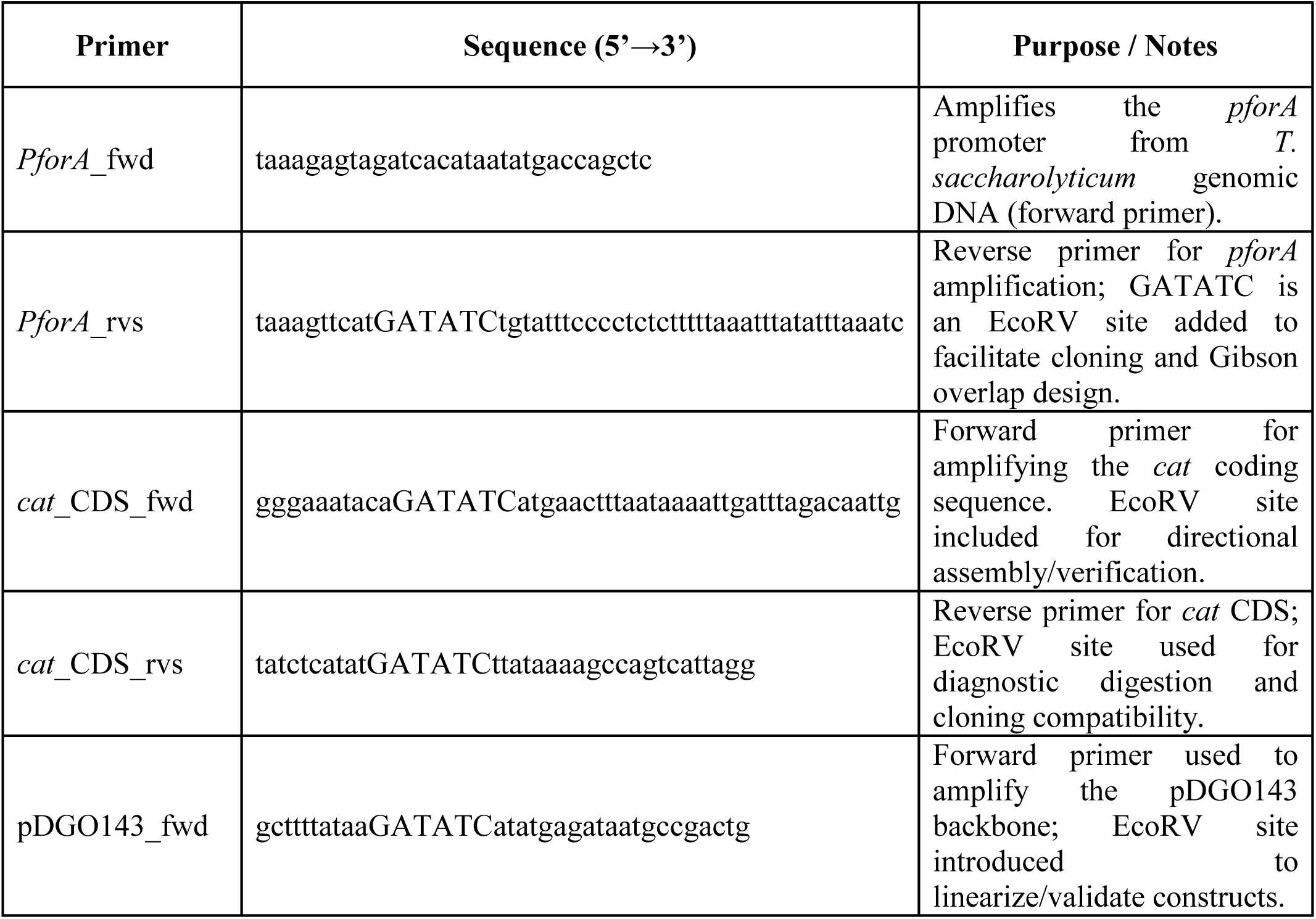
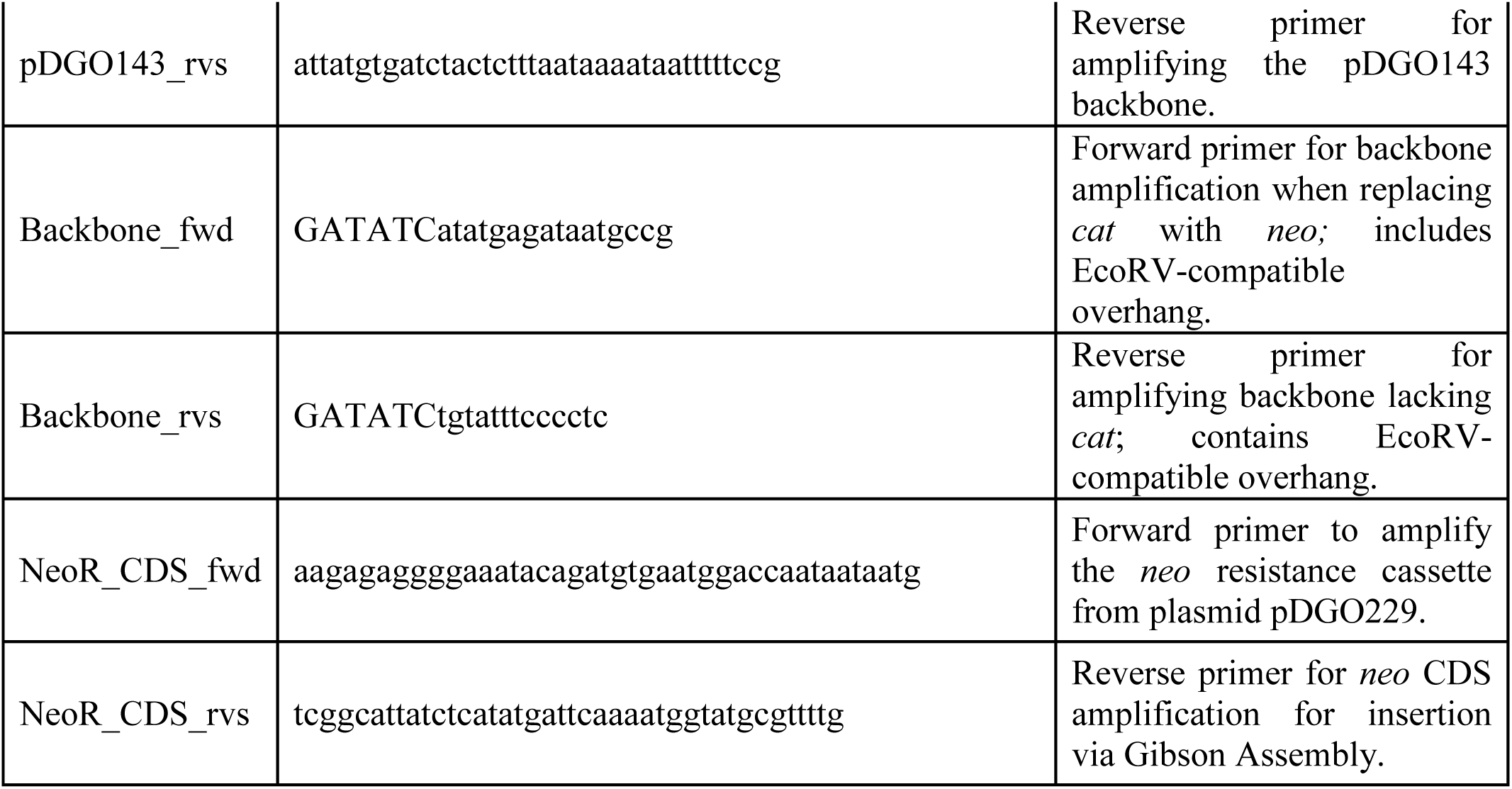
List of primers used for PCR amplification and plasmid construction.

**Table 4.**
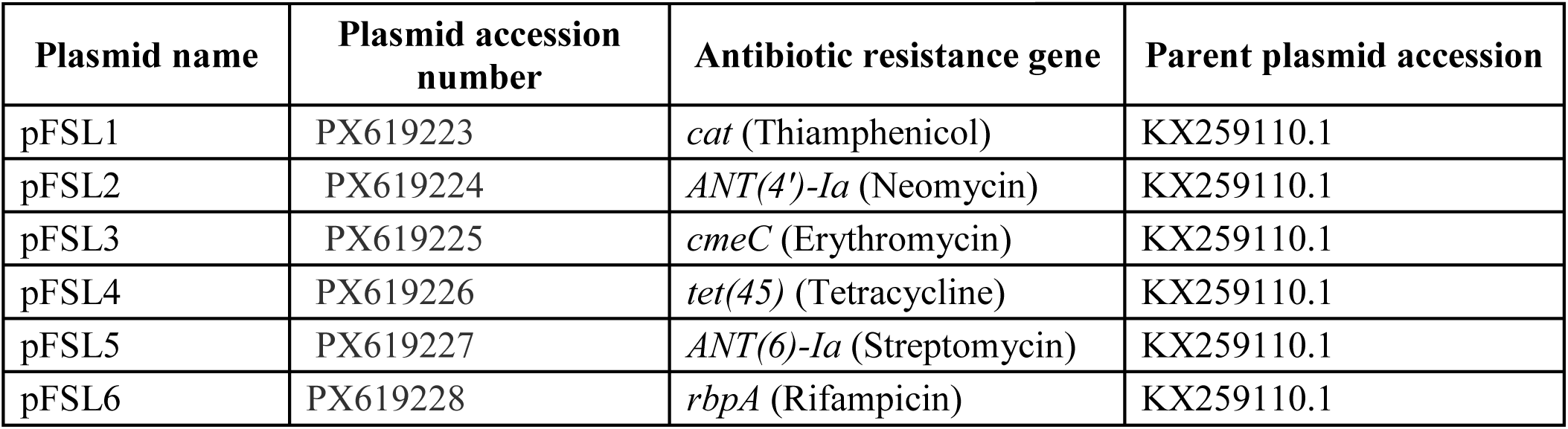
Plasmids constructed and used in this study. Listed are the plasmid names, their GenBank accession numbers, the corresponding antibiotic resistance markers encoded, and the parent plasmid from which each construct was derived.

### 2.4 Transformation of *C. thermocellum* and antibiotic selection

The candidate resistance genes — identified *in silico* as likely to confer resistance to tetracycline, rifampicin, erythromycin, and streptomycin — were synthesized and cloned into plasmid backbones previously constructed for *C. thermocellum*. Transformation assays were then performed to introduce these recombinant plasmids into *C. thermocellum* cells using the standard protocol previously developed by our group (Olson and Lynd, 2012), with modifications optimized for the strains and conditions used in this study. Briefly, cells were grown in a two-step inoculation scheme consisting of 5-mL pre-cultures followed by a 45-mL culture incubated at 55 °C until reaching an OD₆₀₀ of 0.4–0.6. All subsequent handling, including centrifugation, washing, and cuvette preparation, was conducted inside an anaerobic chamber. Cell pellets were washed three times with ultrapure water and resuspended to a final volume of 250 μL; 25 μL of this suspension were mixed on ice with ∼0.5 μg plasmid DNA in pre-chilled 1-mm electroporation cuvettes. Electroporation was performed using a square-wave pulse (1.5 kV, 1.5 ms). Following the pulse, cells were immediately recovered in 5 mL of pre-warmed CTFUD medium and incubated at 50 °C for 18 h before plating onto CTFUD agar containing the appropriate antibiotic for selection.

For each transformation, pre-inocula of *C. thermocellum* were established by adding 5 mL of CTFUD medium to three 15 mL Falcon tubes (or Snapcap tubes) and inoculating with 100, 10, or 1 μL of bacterial stock. Cultures were incubated anaerobically at 55 °C for approximately 18 h. In parallel, 45 mL of CTFUD medium was also incubated overnight at 55 °C to be used for culture expansion. On the following day, 3 mL of an actively growing pre-inoculum was transferred into the 45 mL of pre-warmed CTFUD and incubated anaerobically until the culture reached an OD₆₀₀ between 0.4 and 0.6. Cells were then harvested by centrifugation at 6,700 × g for 15 min and washed three times with sterile ultrapure water under anaerobic conditions. After the final wash, the pellet was gently resuspended in the residual water to a final volume of 250 μL, yielding electrocompetent cells.

Aliquots of 25 μL of these competent cells were mixed with approximately 0.5 μg of plasmid DNA and transferred to pre-chilled 1 mm electroporation cuvettes. Electroporation was performed using a single square-wave pulse at 1,500 V for 1.5 ms. Immediately after pulsing, cells were recovered in 5 mL of pre-warmed CTFUD medium and incubated anaerobically at 50 °C for 18 h.

Recovered cultures were plated onto CTFUD agar containing the corresponding antibiotic for selection. Transformations were carried out using plasmids bearing genes that conferred resistance to tetracycline, rifampicin, or erythromycin. A non-selective control (transformed cells plated without antibiotic) was included in parallel to ensure that any lack of growth was due to antibiotic pressure rather than failed transformation.

The streptomycin-resistance gene was not tested experimentally, as the wild-type *C. thermocellum* strain exhibited a high intrinsic tolerance to streptomycin in previous MIC assays, suggesting a natural resistance to this compound under the tested conditions. Transformants obtained under selective conditions were further evaluated by minimum inhibitory concentration (MIC) assays using liquid CTFUD medium at 48 °C and 60 °C, under the same parameters applied to the wild-type strain, to assess the temperature-dependent expression and stability of each resistance marker.

## 3. Results and Discussion

### 3.1 Distribution and selection of candidate genes

The *in silico* screening identified 1,115 antibiotic resistance genes (ARGs) distributed across 400 thermophilic bacterial genomes. These were classified into 26 drug classes, encompassing 70 Antibiotic Resistance Ontology (ARO) terms in the CARD database (Table 5). Among these, interest resistance determinants were most frequently associated with rifamycin (rifampicin, 8 genes), aminoglycoside (streptomycin, gentamicin, spectinomycin and neomycin, 11 genes), tetracycline (47 genes), macrolide (erythromycin, 11 genes), and phenicol (thiamphenicol, 12 genes) drug classes (Figure 2).

**Figure 2.**
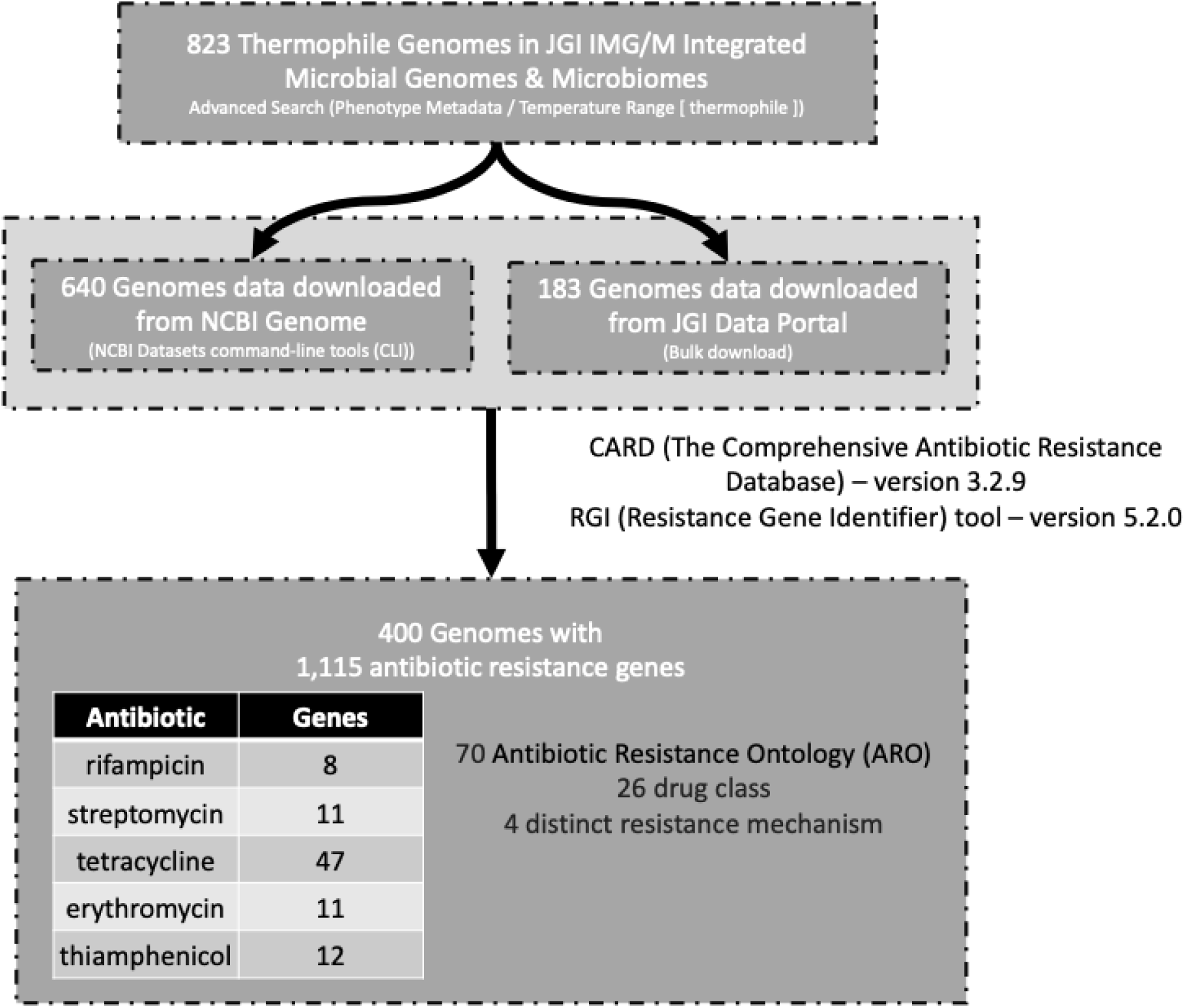
Bioinformatics workflow for thermophile genomes and antibiotic resistance genes identification. A total of 823 thermophilic genomes were retrieved and processed using CARD/RGI, resulting in the detection of 1,115 antibiotic resistance genes across 400 genomes. From these, four antibiotics of interest were selected based on our previously-chosen antibiotics (rifampicin, streptomycin, tetracycline, and erythromycin), leading to 8, 11, 47, and 11 resistance genes identified, respectively.

**Table 5.**
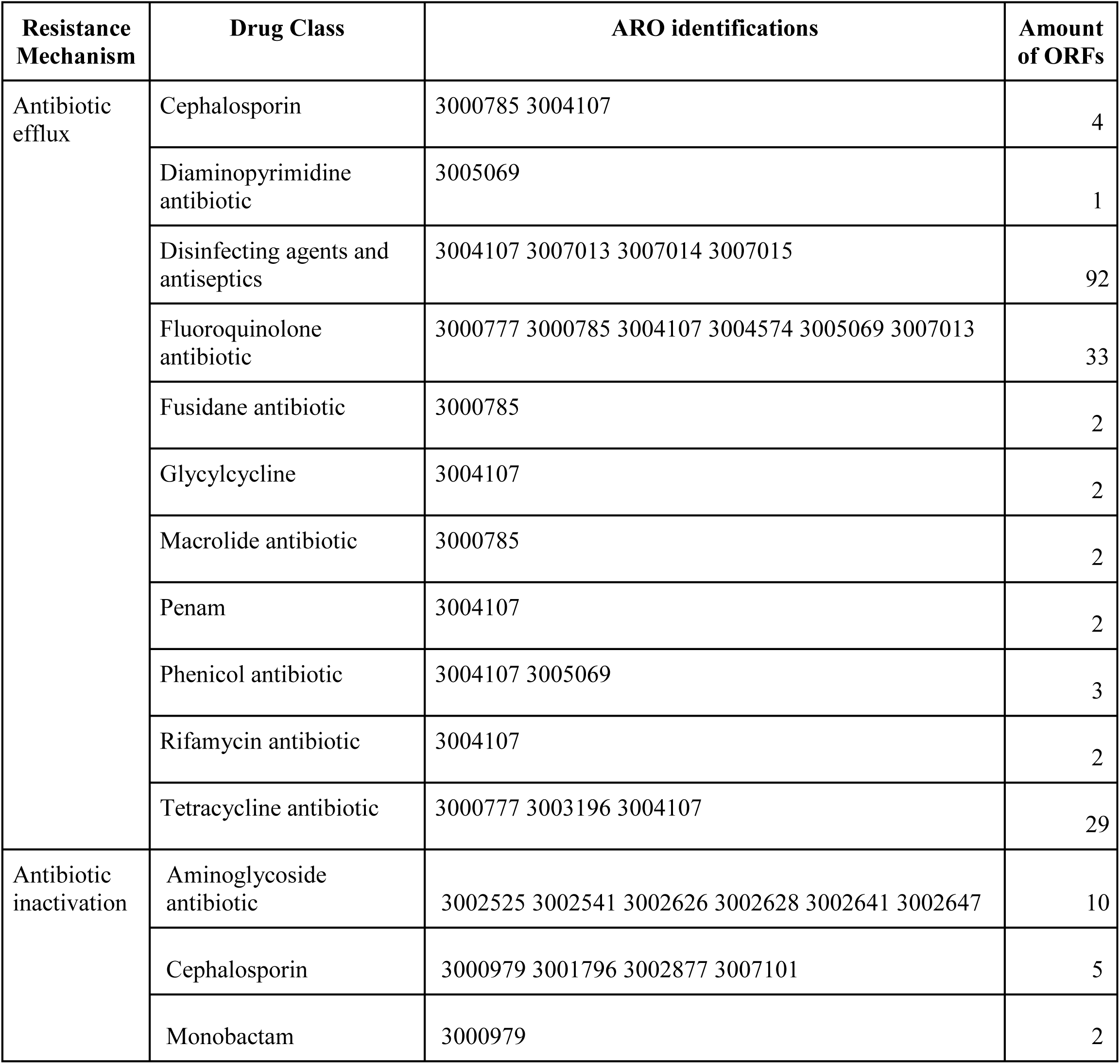
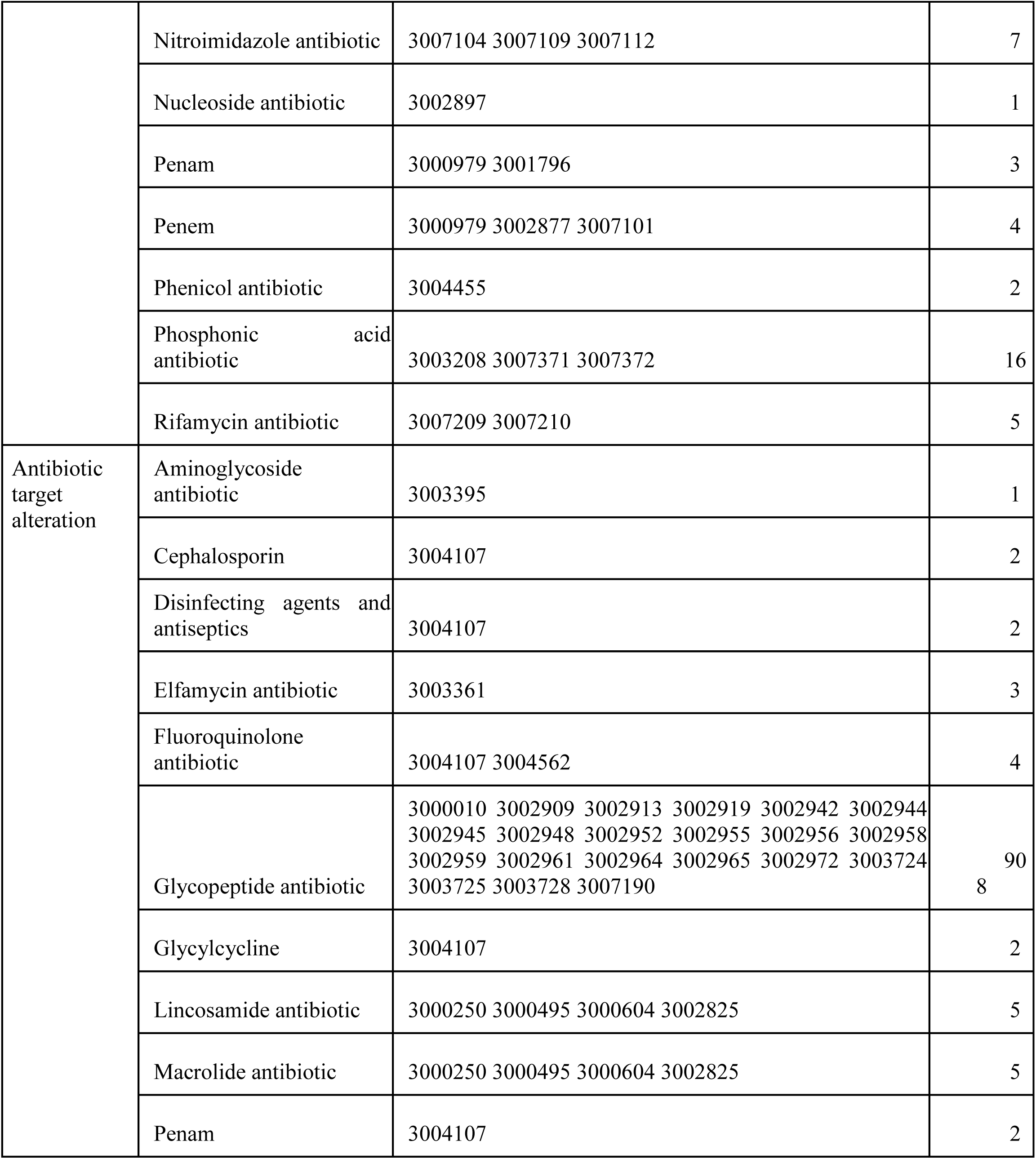
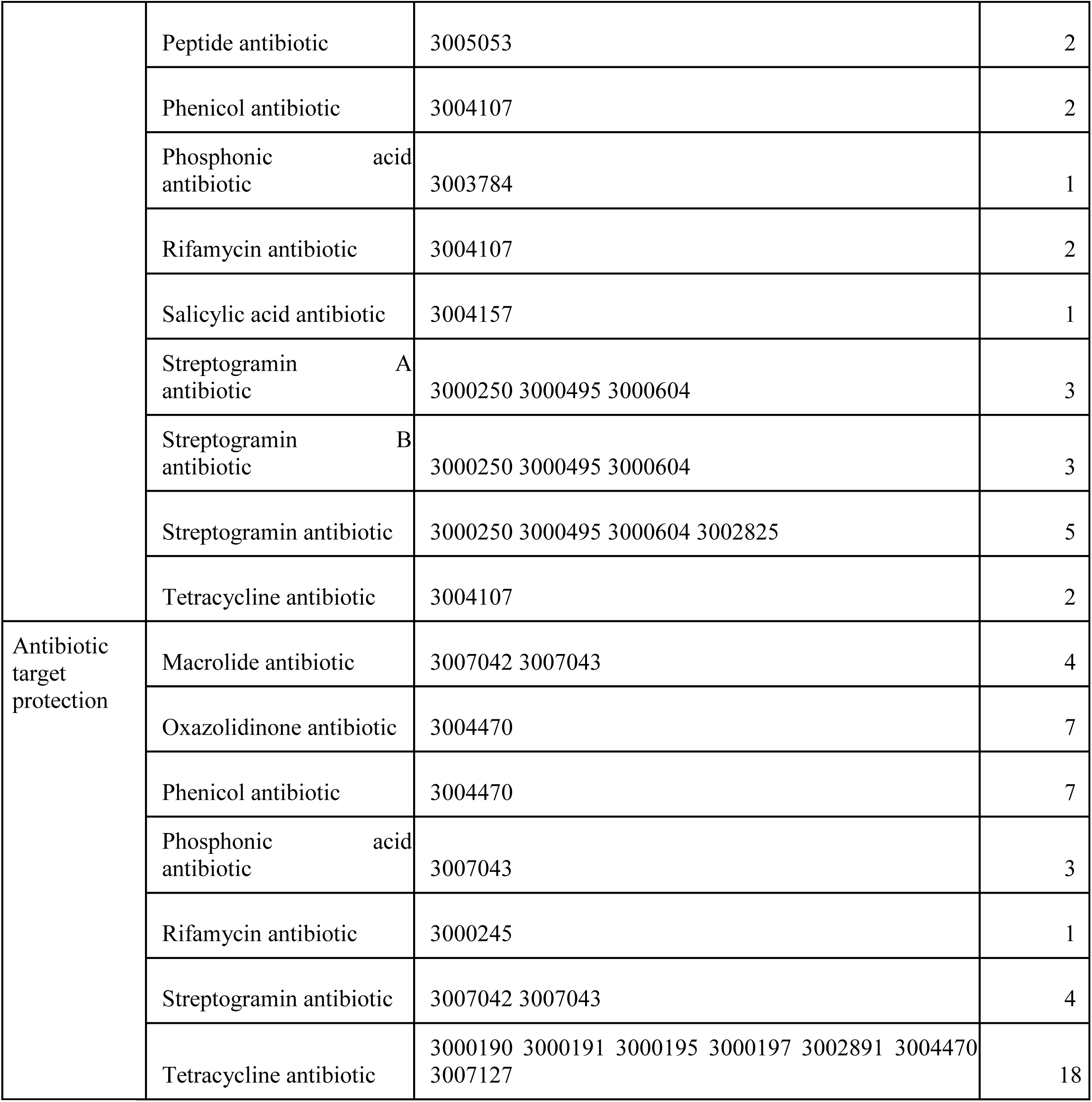
Summary of antibiotic resistance genes (ARGs) identified in thermophilic bacterial genomes. These were classified into 26 drug classes, encompassing 70 Antibiotic Resistance Ontology (ARO) terms in the CARD database.

From the initial dataset (Supplementary Table 1), subsequent analyses were restricted to ARGs both matching the highest percent identity to CARD ARO reference sequences and corresponding to antibiotic classes that were experimentally evaluated in *C. thermocellum* wild type (WT) — namely rifampicin, streptomycin, spectinomycin, gentamicin, neomycin, erythromycin, tetracycline, and thiamphenicol. Due to practical constraints (available funds and time), only the single top-scoring gene (homologous sequence with the highest percentage identity) from each relevant drug class was chosen for synthetic reconstruction and subsequent laboratory validation. The resulting set of prioritized resistance genes used in downstream analyses is listed in Table 6.

**Table 6.**
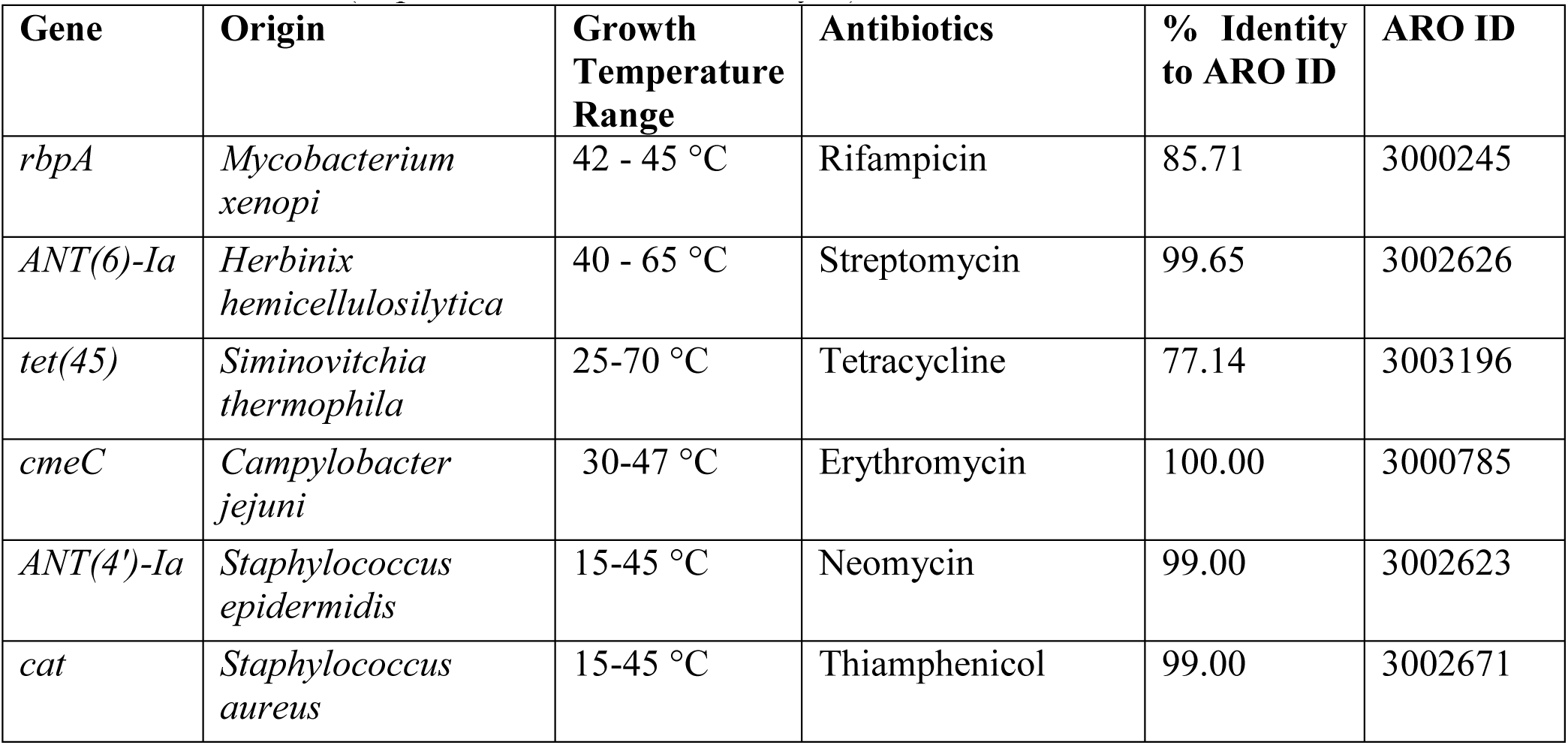
Putative antibiotic resistance genes identified in thermophilic genomes showing the highest similarity to CARD reference sequences. ARO ID (Antibiotic Resistance Ontology ID) designates the controlled vocabulary used by CARD to categorize antimicrobial resistance determinants and enables direct reference to curated entries in the CARD database (https://card.mcmaster.ca/analyze).

This selection resulted in four primary candidates: *rbpA* (rifampicin), *ANT(6)-Ia* (streptomycin), *tet(45)* (tetracycline), and *cmeC* (erythromycin), each representing the highest-identity match to its respective CARD ARO reference sequence. In addition, two previously validated markers, *cat* (thiamphenicol) and *ANT(4’)-Ia* (neomycin), were incorporated as functional controls due to their established activity in thermophilic *Clostridium* systems.

The use of aminoglycoside resistance genes in *C. thermocellum* has been problematic due to low selection range (i.e. only a small difference between the MIC with or without the antibiotic resistance gene) and high variability between laboratories. Natively, *C. thermocellum* is highly resistant to kanamycin (MIC > 1000 ug/ml), however it is slightly more sensitive to neomycin (10-300 ug/ml), and this is the preferred selection system for several recent examples (Lanahan et al., 2022; Olson et al., 2010; Olson, 2011; Walker et al., 2020). Due to this limited selection range, we have often found it necessary to plate neomycin selections on several concentrations of antibiotic to identify a concentration that could distinguish true transformants from background growth.

By contrast, the chloramphenicol acetyltransferase (*cat*) marker, which confers resistance to chloramphenicol and the more thermostable analog thiamphenicol, is well established in *C. thermocellum* genetics and has been used repeatedly as a selection marker (Riley et al., 2019; Tripathi et al., 2010). Importantly, c*at*-based selection using thiamphenicol is preferred for *C. thermocellum* applications due to its high stability and large selection range.

Together, the integration of (i) genome-wide screening, (ii) identity-based ranking of ARGs, (iii) compatibility with antibiotics already evaluated in WT *C. thermocellum*, and (iv) corroboration with existing functional literature resulted in the selection of the most promising candidates for thermophilic selection marker development. These genes, together with their source organisms, temperature ranges, ARO identities, and homology values, are summarized in Table 6.

### 3.2 Antibiotic Sensitivity Testing

The MIC assays revealed clear distinctions between the intrinsic susceptibility profile of wild-type *Clostridium thermocellum* and the phenotypes of transformants carrying heterologous resistance genes (Figure 3, Figure 4). In the wild-type strain, aminoglycosides (neomycin, streptomycin and spectinomycin) produced high MIC values: neomycin showed MICs of ∼32 µg/mL at both 48 °C and 60 °C, while streptomycin and spectinomycin exceeded the upper limit of the assay (>512 µg/mL) (Figure 3), and these two antibiotics were excluded from further consideration.

**Figure 3.**
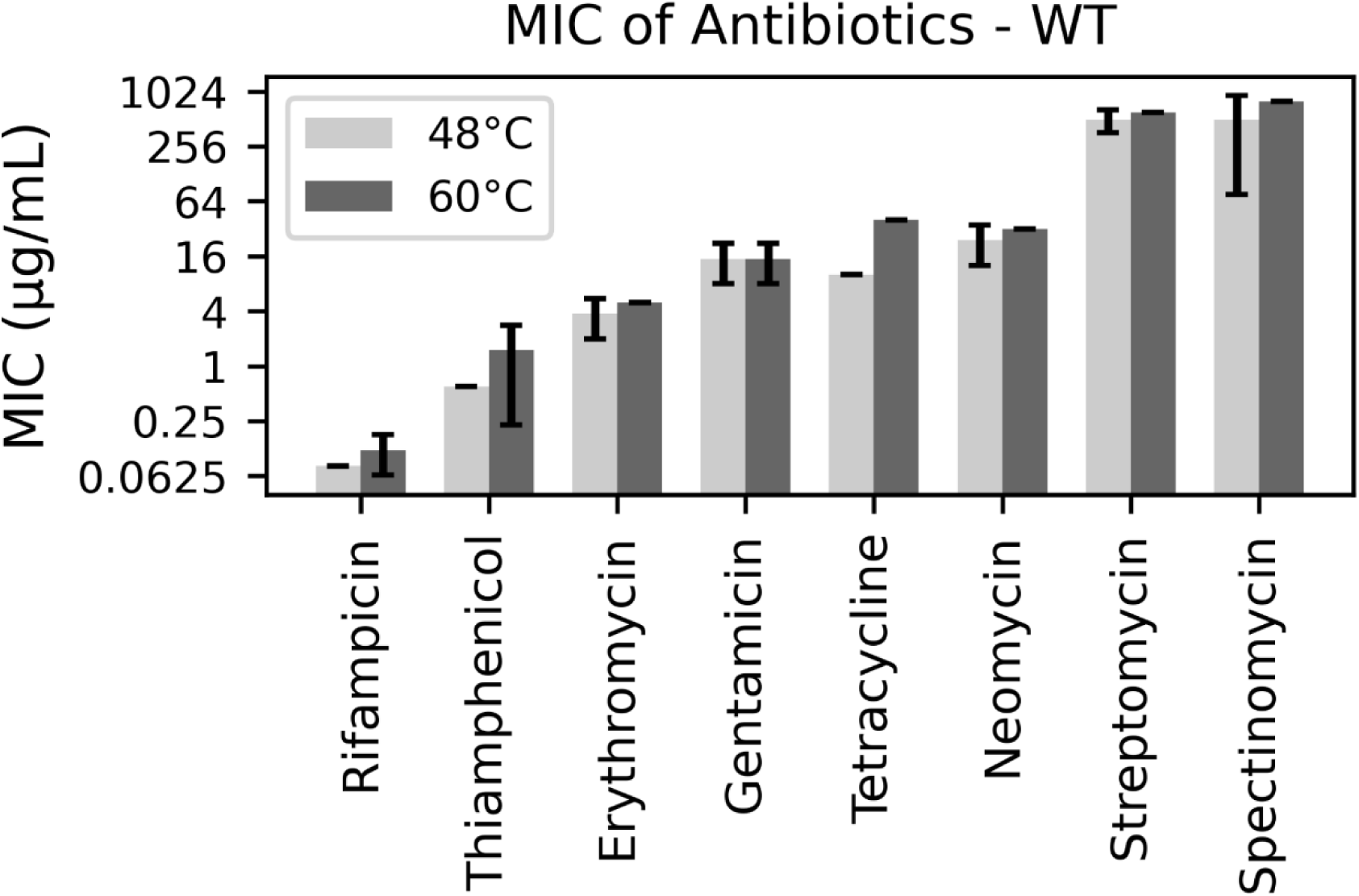
Minimum Inhibitory Concentration (MIC) of antibiotics for the wild-type (WT) strain at two different temperatures. MIC values were determined after 10 days post-inoculation. Light gray bars represent cultures incubated at 48 °C and dark gray bars represent cultures incubated at 60 °C. Error bars indicate the standard deviation (SD) calculated from three independent biological replicates for each condition.

**Figure 4.**
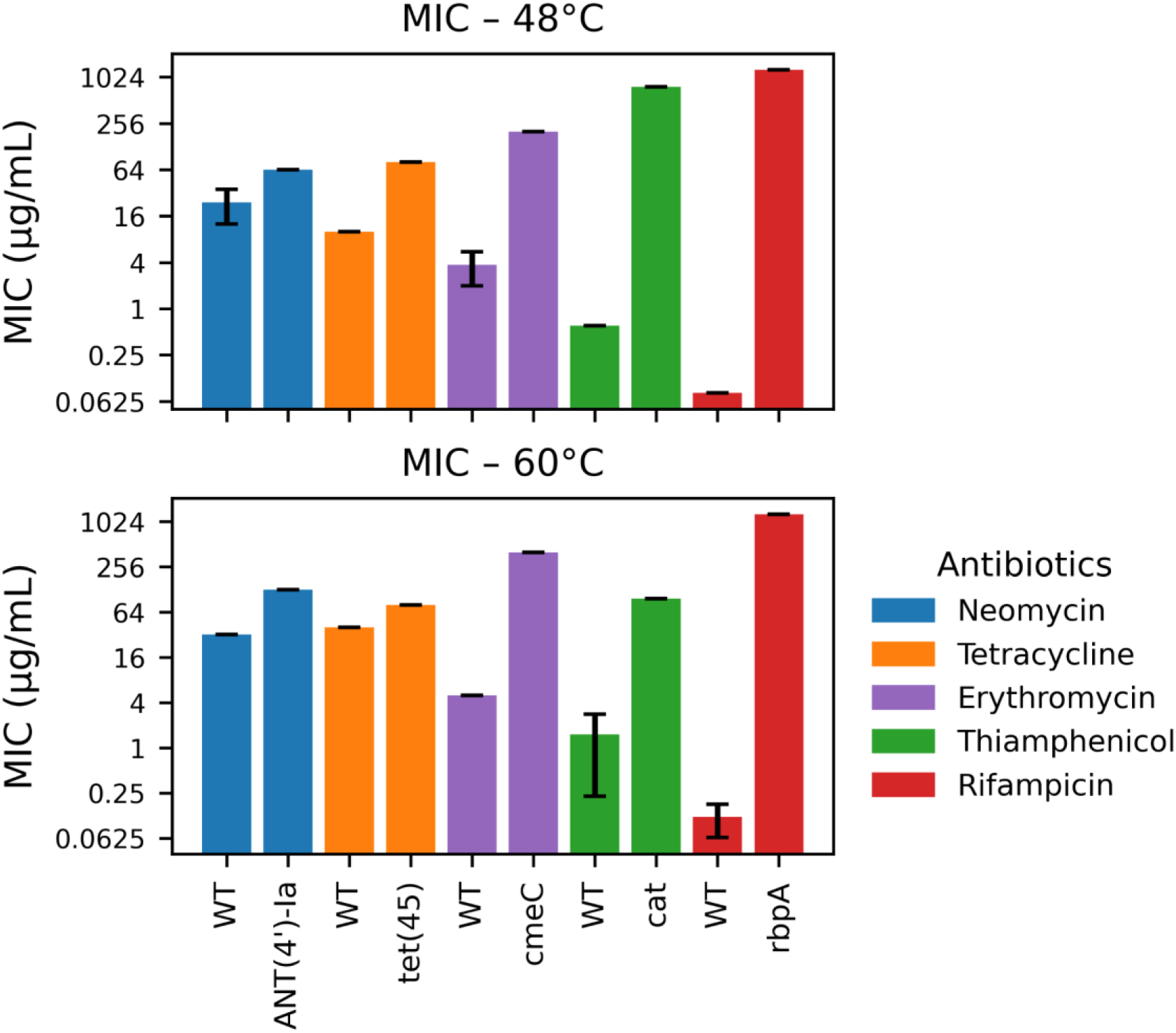
Minimum Inhibitory Concentration (MIC) of five antibiotics for the wild-type (WT) and gene-modified strains at 48 °C (top panel) and 60 °C (bottom panel). MIC values were determined 10 days post-inoculation (10 DPI). Each antibiotic is represented in a different color (blue: Neomycin, green: Thiamphenicol, red: Rifampicin, purple: Erythromycin, Tetracycline: orange). Error bars indicate the standard deviation (SD) calculated from three independent biological replicates for each condition.

By contrast, rifampicin produced the lowest MICs in WT (0.08 µg/mL at 48 °C and 0.16 µg/mL at 60 °C). WT tetracycline MICs were 10 µg/mL at 48 °C and 40 µg/mL at 60°C; tiamphenicol MICs were 0.6 µg/mL (48 °C) and 2.4 µg/mL (60 °C); erythromycin MICs in WT were 2.5–5 µg/mL across temperatures (Figure 3). These baseline values show modest increases in MIC at 60 °C for several antibiotics, suggesting temperature can alter apparent susceptibility (via drug stability, uptake, or target conformation) even in the absence of acquired resistance.

Expression of bioinformatically selected resistance genes produced large, antibiotic-specific increases in MIC (Figure 4). Although the *ANT(4′)-Ia* transformant increased neomycin MIC from 32 µg/mL (WT) to 64 µg/mL at 48 °C and to 128 µg/mL at 60 °C (2× and 4× increases, respectively), this fold-change yields a relatively narrow selection window compared with what is typically considered robust in genetic selection workflows. In practical terms, selection markers that provide a larger dynamic window (often on the order of ∼10-fold or greater) are preferred because they reduce the risk of survival of non-transformed cells and minimize selection of spontaneous or low-level resistant mutants under selective conditions (Drlica, 2003; Gullberg et al., 2011; Guo et al., 2021). Consequently, the modest 2–4× increase observed for *ANT(4′)-Ia*/neomycin suggests a limited selection margin in *C. thermocellum* under our conditions, and therefore neomycin—at least with the current gene/construct—may not be an ideal primary selectable marker for this host.

The transformant bearing *tet(45)* increased the tetracycline MIC from 10 µg/mL to 80 µg/mL at 48 °C (an 8-fold rise) and maintained an elevated MIC of 80 µg/mL at 60 °C (versus 40 µg/mL for the wild-type), supporting the annotation of *tet(45)* as a tetracycline efflux pump (CARD: ARO:3003196) originally described from *Bhargavaea cecembensis* (You et al., 2013). Efflux-mediated resistance has long been recognized as the predominant mechanism underpinning tetracycline resistance—accounting for ∼60 % or more of all tet-related determinants (Blake et al., 2025; Grossman, 2016). The substantial increase at 48°C implies that the efflux system functions efficiently in the thermophilic host under these conditions. However, the narrower fold-difference at 60 °C (2-fold: 40 → 80 µg/mL) highlights two potential caveats: first, the wild-type baseline MIC itself rose at the elevated temperature, reducing the dynamic window available for selection; second, the efflux pump’s relative benefit did not scale further with temperature, which may indicate that membrane transport, pump stability, or tetracycline influx/egress dynamics are altered in the thermophilic context. From a selectable marker perspective, while *tet(45)* clearly confers functional resistance in *C. thermocellum*, the narrow selection range may make it difficult to use in practice.

The transformant carrying *cmeC* showed a striking increase in erythromycin MIC compared with the wild type (from 2.5–5 µg/mL in WT to 200 µg/mL at 48 °C and 400 µg/mL at 60 °C, corresponding to ∼40–80× increases) (Figure 4). Erythromycin and other macrolides act by binding to a conserved site in the 50S ribosomal subunit, partially occluding the nascent peptide exit tunnel and preventing normal peptide elongation (Dinos, 2017). In many bacteria, resistance to macrolides is mediated by efflux-associated envelope proteins, including *CmeC* homologs (Alav et al., 2021; Gibreel et al., 2007; Su et al., 2017). Although the precise function of *cmeC* when expressed in *C. thermocellum* is not fully defined, the strong increase in erythromycin MIC indicates that the protein can provide effective protection in this thermophilic host.

Temperature effects on efflux activity are context-dependent and species-specific; some studies report decreased efflux at very high temperatures while others find increased substrate transport owing to altered membrane properties (Blair and Piddock, 2016). Importantly for marker development, the very large selection window created by *cmeC* (tens to hundreds of fold) provides a strong discriminatory margin between WT and transformants, making *cmeC* a promising candidate for macrolide-based selection in *C. thermocellum*.

The transformant carrying *cat* conferred one of the largest increases in thiamphenicol resistance observed in this study: MIC rose from 0.6 µg/mL in the wild-type to ∼768 µg/mL at 48 °C (>1000× increase), but decreased markedly to 96 µg/mL at 60 °C (an ∼8-fold reduction relative to 48 °C) (Figure 4). This pattern indicates that *cat* expression can provide a very strong resistance phenotype in *C. thermocellum*, but that the effective activity of the enzyme or the intracellular conditions required for its optimal function is impaired at the highest growth temperature tested.

Chloramphenicol acetyltransferase genes (*cats*) mediate resistance by acetylating chloramphenicol and related phenicols (including thiamphenicol), and *cat* genes have a long history as convenient selectable markers in a wide range of bacteria and organelles, because enzymatic inactivation produces a high selection window when the enzyme is active (Xu et al., 2013). Once acetylated, the modified phenicol can no longer bind the peptidyl-transferase center of the 50S ribosomal subunit, resulting in complete loss of inhibitory activity. The acetylation also increases the hydrophobicity of the molecule, which promotes passive diffusion or nonspecific efflux out of the cell, further reducing the intracellular concentration of the active drug (Murray and Shaw, 1997; Schwarz et al., 2004). These combined effects make *cat*-mediated inactivation highly efficient and explain why *cat* often provides a broad and robust selection window in thermophiles when the enzyme remains active.

Importantly for thermophilic applications, both natural *cat* variants with elevated thermal tolerance and engineered thermostable *cat*s have been described: early selection experiments isolated thermostable *cat* mutants with enhanced activity above 50 °C (Turner et al., 1992), and more recent work has used protein engineering to produce *cat* variants that retain activity at elevated temperatures and have been repurposed for thermophilic bioproduction workflows (Seo et al., 2019). These studies demonstrate that *cat* can be adapted for high-temperature use, but they also underscore that not all *cat* alleles are intrinsically thermostable. In our case, the pronounced drop in MIC at 60 °C suggests partial thermolability of the *cat* allele used (or altered intracellular cofactor availability or folding/assembly at higher temperature), meaning that the current *cat* construct may require thermostabilizing mutations (or a different *cat* ortholog) to function reliably as a selectable marker at industrially relevant thermophilic temperatures (Seo et al., 2019). Finally, practical precedent exists for using related phenicol resistance markers in non-model bacteria, supporting the general strategy of phenicol selection in diverse hosts while highlighting the need to verify marker performance under the intended growth conditions (Groom et al., 2016; Sanford and Woolston, 2022).

The most striking phenotype observed was conferred by *rbpA*: transformants expressing this gene exhibited rifampicin MICs >1280 µg/mL at both 48 °C and 60 °C, corresponding to an increase of more than 10,000-fold relative to the wild-type baseline (0.08–0.16 µg/mL) (Figure 4). This strong rifampicin resistance phenotype aligns with the established function of RbpA homologs, which bind RNA polymerase and stabilize the initiation complex in the presence of rifampicin, reducing the drug’s inhibitory effect on transcription initiation (Hubin et al., 2017; Newell et al., 2006). Structural studies of mycobacterial transcription complexes show that *rbpA* binds near the upstream edge of the −10 promoter element and can stabilize the open complex, a mechanism that plausibly explains how *rbpA* protects RNAP from rifampicin binding without requiring *rpoB* target mutations (Hubin et al., 2017). The magnitude and thermal robustness of the *rbpA* phenotype make rifampicin selection particularly attractive for *C. thermocellum*: a very low WT MIC provides a broad dynamic window for selection, and the *rbpA* gene confers strong protection at operational temperatures used for this organism.

Rifampicin resistance in many bacteria commonly arises through target-site mutations in RNA polymerase, but such mutations can impose pleiotropic fitness costs and alter global transcription patterns (Cutugno et al., 2020). Using *rbpA* as a resistance determinant therefore offers practical advantages: (i) it provides very high-level resistance without requiring chromosomal mutations that may be deleterious, and (ii) its mechanism is based on protein-mediated protection of RNA polymerase rather than permanent target alteration, which may preserve physiological function more faithfully. Nevertheless, *rbpA* is a transcription-associated factor, and its heterologous expression could influence transcriptional programs or impose fitness costs in the absence of selection. Prior work in actinobacteria indicates that *rbpA* can modulate rRNA promoter activity and global transcription (Newell et al., 2006). Rifampicin has long been used as a genetic marker because strong resistance phenotypes are easy to select and detect, but the stability and physiological neutrality of resistance must be confirmed for each host (Glandorf et al., 1992). Taken together, our data indicate that *rbpA* provides an exceptionally large and thermally stable selection window in *C. thermocellum* and is therefore a strong candidate for rifampicin-based selection.

Together, these results demonstrate that thermophilic hosts can support a broad range of antibiotic resistance mechanisms, and that most antibiotic–resistance phenotypes in *C. thermocellum* remain largely stable across the two operational temperatures tested (48°C and 60 °C). Only a subset of determinants, most notably *cat*, which shows reduced thiamphenicol protection at 60 °C, and *cmeC*, which exhibits a modest temperature-dependent shift, display measurable thermal sensitivity. In contrast, neomycin, tetracycline, and rifampicin resistance levels are essentially unaffected by temperature.

Among all tested determinants, *rbpA* emerges as an outstanding candidate marker for thermophilic selection, combining the intrinsically low WT rifampicin MIC with extremely high, temperature-stable transformed MICs. This results in the widest and most reliable selection window observed in this study, making *rbpA*-mediated rifampicin resistance particularly powerful for high-temperature genetic engineering applications.

Beyond validating individual markers, the study supports a generalizable workflow for discovering and applying resistance determinants in thermophiles: mining genome databases, selecting candidates based on similarity to curated references, assembling constructs through cloning or gene synthesis, and testing function directly in the thermophilic host. Under this framework, *cat* and *cmeC* also provide strong options for phenicol and macrolide selection, respectively.

Overall, the data indicate that thermophilic antibiotic selection is both feasible and robust, enabling flexible genetic manipulation of *C. thermocellum* and establishing a foundation for expanding the genetic toolkit available for high-temperature industrial microorganisms.

## Supplementary Information

**Supplementary Table 1.** Complete data set of the 1,115 antibiotic resistance genes (ARGs) identified in this study, including ORF identification, best hit ARO, hit identity, model type, drug class, resistance mechanism, and predict protein sequence.

### DNA sequences of all antibiotic resistance genes

>cat_gene

atgaactttaataaaattgatttagacaattggaagagaaaagagatatttaatcattatttgaaccaacaaacgacttttagtataacca cagaaattgatattagtgttttataccgaaacataaaacaagaaggatataaattttaccctgcatttattttcttagtgacaagggtgata aactcaaatacagcttttagaactggttacaatagcgacggagagttaggttattgggataagttagagccactttatacaatttttgatg gtgtatctaaaacattctctggtatttggactcctgtaaagaatgacttcaaagagttttatgatttatacctttctgatgtagagaaatataa tggttcggggaaattgtttcccaaaacacctatacctgaaaatgctttttctctttctattattccatggacttcatttactgggtttaacttaa atatcaataataatagtaattaccttctacccattattacagcaggaaaattcattaataaaggtaattcaatatatttaccgctatctttaca ggtacatcattctgtttgtgatggttatcatgcaggattgtttatgaactctattcaggaattgtcagataggcctaatgactggcttttata a

>ANT(4’)-Ia_gene gtgaatggaccaataataatgactagagaagaaagaatgaagattgttcatgaaattaaggaacgaatattggataaatatggggat gatgttaaggctattggtgtttatggctctcttggtcgtcagactgatgggccctattcggatattgagatgatgtgtgtcatgtcaacag aggaagcagagttcagccatgaatggacaaccggtgagtggaaggtggaagtgaattttgatagcgaagagattctactagattat gcatctcaggtggaatcagattggccgcttacacatggtcaatttttctctattttgccgatttatgattcaggtggatacttagagaaagt gtatcaaactgctaaatcggtagaagcccaaacgttccacgatgcgatttgtgcccttatcgtagaagagctgtttgaatatgcaggc aaatggcgtaatattcgtgtgcaaggaccgacaacatttctaccatccttgactgtacaggtagcaatggcaggtgccatgttgattgg tctgcatcatcgcatctgttatacgacgagcgcttcggtcttaactgaagcagttaagcaatcagatcttccttcaggttatgaccatctg tgccagttcgtaatgtctggtcaactttccgactctgagaaacttctggaatcgctagagaatttctggaatgggattcaggagtggac agaacgacacggatatatagtggatgtgtcaaaacgcataccattttga

>*cmeC*_gene

atgaataaaattatttcaatttcagcaattgcatcatttacattattaatttcagcatgttcattatcacctaatttaaatattcctgaagcaaat tattcaattgataataaattaggagcattatcatgggaaaaagaaaataattcatcaattacaaaaaattggtggaaagattttgatgatg aaaatttaaataaagttgttgatttagcattaaaaaataataatgatttaaaattagcatttattcatatggaacaagcagcagcacaatta ggaattgatttttcatcattattacctaaatttgatggatcagcatcaggatcaagagcaaaaacagcaattaatgcaccttcaaataga acaggagaagtttcatatggaaatgattttaaaatgggattaaatttatcatatgaaattgatttatggggaaaatatagagatacatata gagcatcaaaatcatcatttaaagcatcagaatatgattatgaagcagcaagattatcagttatttcaaatacagttcaaacatattttaat ttagttaatgcatatgaaaatgaaaatgcattaaaagaagcatatgaatcagcaaaagaaatttatagaattaatgatgaaaaatttcaa gttggagcagttggagaatatgaattagcacaagcaagagcaaatttagaatcaatggcattacaatataatgaagcaaaattaaata

aagaaaattatttaaaagcattaaaaattttaacatcaaatgatttaaatgatattttatataaaaatcaatcatatcaagtttttaatttaaaa gaatttgatattcctacaggaatttcatcaacaattttattacaaagacctgatattggatcatcattagaaaaattaacacaacaaaattat ttagttggagttgcaagaacagcatttttaccttcattatcattaacaggattattaggatttgaatcaggagatttagatacattagttaaa ggaggatcaaaaacatggaatattggaggaaattttacattacctatttttcattggggagaaatttatcaaaatgttaatttagcaaaatt aaataaagatgaagcatttgttaattatcaaaatacattaattacagcatttggagaaattagatatgcattagttgcaagaaaaacaatt agattacaatatgataatgcacaagcatcagaacaatcatataaaagaatttatgaaattgcaaaagaaagatatgatattggagaaat gtcattacaagattatttagaagcaagacaaaattggttaaatgcagcagttgcatttaataatacaaaatattcatatgcaaattcaatta ttgatgttattaaagcatttggaggaggatttgaacaatcagaagatacatcaaaaaatattaaagaagaatcaaaaaatttagatatgt catttagagaatga

>*tet(45)*_gene

atgaatacatcatattcacaatcaaatttaagacataatcaaattttaatttggttatgtattttatcatttttttcagttttaaatgaaatggtttt aaatgtttcattacctgatattgcaaatgattttaataaacctcctgcatcaacaaattgggttaatacagcatttatgttaacattttcaattg gaacagcagtttatggaaaattatcagatcaattaggaattaaaagattattattatttggaattattattaattgttttggatcagttattgga tttgttggacattcatttttttcattattaattatggcaagatttattcaaggagcaggagcagcagcatttcctgcattagttatggttgttgt tgcaagatatattcctaaagaaaatagaggaaaagcatttggattaattggatcaattgttgcaatgggagaaggagttggacctgca attggaggaatgattgcacattatattcattggtcatatttattattaattcctatgattacaattattacagttccttttttaatgaaattattaaa aaaagaagttagaattaaaggacattttgatattaaaggaattattttaatgtcagttggaattgttttttttatgttatttacaacatcatattc aatttcatttttaattgtttcagttttatcatttttaatttttgttaaacatattagaaaagttacagatccttttgttgatcctggattaggaaaaa atattccttttatgattggagttttatgtggaggaattatttttggaacagttgcaggatttgtttcaatggttccttatatgatgaaagatgtt catcaattatcaacagcagaaattggatcagttattatttttcctggaacaatgtcagttattatttttggatatattggaggaattttagttga tagaagaggacctttatatgttttaaatattggagttacatttttatcagtttcatttttaacagcatcatttttattagaaacaacatcatggttt atgacaattattattgtttttgttttaggaggattatcatttacaaaaacagttatttcaacaattgtttcatcatcattaaaacaacaagaagc aggagcaggaatgtcattattaaattttacatcatttttatcagaaggaacaggaattgcaattgttggaggattattatcaattcctttatt agatcaaagattattacctatggaagttgatcaatcaacatatttatattcaaatttattattattattttcaggaattattgttatttcatggtta gttacattaaatgtttataaacattcacaatga

>*rbpA*_gene

atggcagatagagttttaagaggatcaagattaggagcagtttcatatgaaacagatagaaatcatgatttagcacctagacaaattgc aagatatagaacagaaaatggagaagtttttgaagttccttttgcagatgatgcagaaattcctggaacatggttatgtagaaatggaa tggaaggagttttaattgaaggagatcaacctgaacctaaaaaagttaaacctcctagaacacattgggatatgttattagaaagaag aacaattgaagaattagatgaattattaaaagaaagattagaaattattagacaaagaagaagaggatcatga

## Author contributions

Fernanda - Formal analysis; investigation; writing. Renato Vicentini - Data curation; formal analysis. Luana Walravens Bergamo - Analysis; writing.

Edson Kim - Data curation; writing Nandhini Ashok – Investigation.

Adam M. Guss – Investigation.

Lee R. Lynd - Funding acquisition; project administration; supervision; resources; writing. Daniel G. Olson - Formal analysis; investigation; writing; supervision; conceptualization.

## References

1. Alav, I., Kobylka, J., Kuth, M.S., Pos, K.M., Picard, M., Blair, J.M.A., Bavro, V.N., 2021. Structure, Assembly, and Function of Tripartite Efflux and Type 1 Secretion Systems in Gram-Negative Bacteria. Chem. Rev. 121, 5479–5596. 10.1021/acs.chemrev.1c00055

2. Alcock, B.P., Huynh, W., Chalil, R., Smith, K.W., Raphenya, A.R., Wlodarski, M.A., Edalatmand, A., Petkau, A., Syed, S.A., Tsang, K.K., Baker, S.J.C., Dave, M., McCarthy, M.C., Mukiri, K.M., Nasir, J.A., Golbon, B., Imtiaz, H., Jiang, X., Kaur, K., Kwong, M., Liang, Z.C., Niu, K.C., Shan, P., Yang, J.Y.J., Gray, K.L., Hoad, G.R., Jia, B., Bhando, T., Carfrae, L.A., Farha, M.A., French, S., Gordzevich, R., Rachwalski, K., Tu, M.M., Bordeleau, E., Dooley, D., Griffiths, E., Zubyk, H.L., Brown, E.D., Maguire, F., Beiko, R.G., Hsiao, W.W.L., Brinkman, F.S.L., Van Domselaar, G., McArthur, A.G., 2023. CARD 2023: expanded curation, support for machine learning, and resistome prediction at the Comprehensive Antibiotic Resistance Database. Nucleic Acids Res. 51, D690–D699. 10.1093/nar/gkac920

3. Blair, J.M.A., Piddock, L.J.V., 2016. How to Measure Export via Bacterial Multidrug Resistance Efflux Pumps. mBio 7, e00840–16. 10.1128/mBio.00840-16

4. Blake, K.S., Xue, Y.-P., Gillespie, V.J., Fishbein, S.R.S., Tolia, N.H., Wencewicz, T.A., Dantas, G., 2025. The tetracycline resistome is shaped by selection for specific resistance mechanisms by each antibiotic generation. Nat. Commun. 16, 1452. 10.1038/s41467-025-56425-5

5. Brouns, S.J.J., Wu, H., Akerboom, J., Turnbull, A.P., De Vos, W.M., Van Der Oost, J., 2005. Engineering a Selectable Marker for Hyperthermophiles. J. Biol. Chem. 280, 11422–11431. 10.1074/jbc.M413623200

6. Buchfink, B., Xie, C., Huson, D.H., 2015. Fast and sensitive protein alignment using DIAMOND. Nat. Methods 12, 59–60. 10.1038/nmeth.3176

7. Canadell, J.G., Schulze, E.D., 2014. Global potential of biospheric carbon management for climate mitigation. Nat. Commun. 5, 5282. 10.1038/ncomms6282

8. Carere, C.R., Sparling, R., Cicek, N., Levin, D.B., 2008. Third Generation Biofuels via Direct Cellulose Fermentation. Int. J. Mol. Sci. 9, 1342–1360. 10.3390/ijms9071342

9. Chen, I.-M.A., Chu, K., Palaniappan, K., Ratner, A., Huang, J., Huntemann, M., Hajek, P., Ritter, S.J., Webb, C., Wu, D., Varghese, N.J., Reddy, T.B.K., Mukherjee, S., Ovchinnikova, G., Nolan, M., Seshadri, R., Roux, S., Visel, A., Woyke, T., Eloe-Fadrosh, E.A., Kyrpides, N.C., Ivanova, N.N., 2023. The IMG/M data management and analysis system v.7: content updates and new features. Nucleic Acids Res. 51, D723–D732. 10.1093/nar/gkac976

10. Cutugno, L., Mc Cafferty, J., Pané-Farré, J., O’Byrne, C., Boyd, A., 2020. rpoB mutations conferring rifampicin-resistance affect growth, stress response and motility in Vibrio vulnificus. Microbiology 166, 1160–1170. 10.1099/mic.0.000991

11. Dinos, G.P., 2017. The macrolide antibiotic renaissance. Br. J. Pharmacol. 174, 2967–2983. 10.1111/bph.13936

12. Drlica, K., 2003. The mutant selection window and antimicrobial resistance. J. Antimicrob. Chemother. 52, 11–17. 10.1093/jac/dkg269

13. Field, J.L., Richard, T.L., Smithwick, E.A.H., Cai, H., Laser, M.S., LeBauer, D.S., Long, S.P., Paustian, K., Qin, Z., Sheehan, J.J., Smith, P., Wang, M.Q., Lynd, L.R., 2020. Robust paths to net greenhouse gas mitigation and negative emissions via advanced biofuels. Proc. Natl. Acad. Sci. 117, 21968–21977. 10.1073/pnas.1920877117

14. Gibreel, A., Wetsch, N.M., Taylor, D.E., 2007. Contribution of the CmeABC Efflux Pump to Macrolide and Tetracycline Resistance in *Campylobacter jejuni*. Antimicrob. Agents Chemother. 51, 3212–3216. 10.1128/AAC.01592-06

15. Glandorf, D.C.M., Brand, I., Bakker, P.A.H.M., Schippers, B., 1992. Stability of rifampicin resistance as a marker for root colonization studies of Pseudomonas putida in the field. Plant Soil 147, 135–142. 10.1007/BF00009379

16. Groom, J., Chung, D., Olson, D.G., Lynd, L.R., Guss, A.M., Westpheling, J., 2016. Promiscuous plasmid replication in thermophiles: Use of a novel hyperthermophilic replicon for genetic manipulation of Clostridium thermocellum at its optimum growth temperature. Metab. Eng. Commun. 3, 30–38. 10.1016/j.meteno.2016.01.004

17. Grossman, T.H., 2016. Tetracycline Antibiotics and Resistance. Cold Spring Harb. Perspect. Med. 6, a025387. 10.1101/cshperspect.a025387

18. Gullberg, E., Cao, S., Berg, O.G., Ilbäck, C., Sandegren, L., Hughes, D., Andersson, D.I., 2011. Selection of Resistant Bacteria at Very Low Antibiotic Concentrations. PLoS Pathog. 7, e1002158. 10.1371/journal.ppat.1002158

19. Guo, C., Fordjour, F.K., Tsai, S.J., Morrell, J.C., Gould, S.J., 2021. Choice of selectable marker affects recombinant protein expression in cells and exosomes. J. Biol. Chem. 297, 100838. 10.1016/j.jbc.2021.100838

20. Haima, P., Bron, S., Venema, G., 1987. The effect of restriction on shotgun cloning and plasmid stability in Bacillus subtilis Marburg. Mol. Gen. Genet. MGG 209, 335–342. 10.1007/BF00329663

21. Hon, S., Holwerda, E.K., Worthen, R.S., Maloney, M.I., Tian, L., Cui, J., Lin, P.P., Lynd, L.R., Olson, D.G., 2018. Expressing the Thermoanaerobacterium saccharolyticum pforA in engineered Clostridium thermocellum improves ethanol production. Biotechnol. Biofuels 11, 242. 10.1186/s13068-018-1245-2

22. Hon, S., Lanahan, A.A., Tian, L., Giannone, R.J., Hettich, R.L., Olson, D.G., Lynd, L.R., 2016. Development of a plasmid-based expression system in Clostridium thermocellum and its use to screen heterologous expression of bifunctional alcohol dehydrogenases (adhEs). Metab. Eng. Commun. 3, 120–129. 10.1016/j.meteno.2016.04.001

23. Hubin, E.A., Fay, A., Xu, C., Bean, J.M., Saecker, R.M., Glickman, M.S., Darst, S.A., Campbell, E.A., 2017. Structure and function of the mycobacterial transcription initiation complex with the essential regulator RbpA. eLife 6, e22520. 10.7554/eLife.22520

24. Kortam, Y.G., Abd El-Rahim, W.M., Khattab, A.E.-N.A., Rebouh, N.Y., Gurina, R.R., Barakat, O.S., Zakaria, M., Moawad, H., 2023. Enhancing the Antibiotic Production by Thermophilic Bacteria Isolated from Hot Spring Waters via Ethyl Methanesulfonate Mutagenesis. Antibiotics 12, 1095. 10.3390/antibiotics12071095

25. Lanahan, A., Zakowicz, K., Tian, L., Olson, D.G., Lynd, L.R., 2022. A Single Nucleotide Change in the *polC* DNA Polymerase III in Clostridium thermocellum Is Sufficient To Create a Hypermutator Phenotype. Appl. Environ. Microbiol. 88, e01531–21. 10.1128/AEM.01531-21

26. Lynd, L.R., Weimer, P.J., Van Zyl, W.H., Pretorius, I.S., 2002. Microbial Cellulose Utilization: Fundamentals and Biotechnology. Microbiol. Mol. Biol. Rev. 66, 506–577. 10.1128/MMBR.66.3.506-577.2002

27. Mai, V., Lorenz, W.W., Wiegel, J., 2006. Transformation of Thermoanaerobacterium sp. strain JW/SL-YS485 with plasmid pIKM1 conferring kanamycin resistance. FEMS Microbiol. Lett. 148, 163–167. 10.1111/j.1574-6968.1997.tb10283.x

28. Murray, I.A., Shaw, W.V., 1997. O-Acetyltransferases for chloramphenicol and other natural products. Antimicrob. Agents Chemother. 41, 1–6. 10.1128/AAC.41.1.1

29. Newell, K.V., Thomas, D.P., Brekasis, D., Paget, M.S.B., 2006. The RNA polymerase-binding protein RbpA confers basal levels of rifampicin resistance on *Streptomyces coelicolor*. Mol. Microbiol. 60, 687–696. 10.1111/j.1365-2958.2006.05116.x

30. Noll, K.M., Vargas, M., 1997. Recent advances in genetic analyses of hyperthermophilic Archaea and Bacteria. Arch. Microbiol. 168, 73–80. 10.1007/s002030050472

31. Okada, B.K., Seyedsayamdost, M.R., 2017. Antibiotic dialogues: induction of silent biosynthetic gene clusters by exogenous small molecules. FEMS Microbiol. Rev. 41, 19–33. 10.1093/femsre/fuw035

32. Olson, D.G., Lynd, L.R., 2012. Transformation of Clostridium Thermocellum by Electroporation, in: Methods in Enzymology. Elsevier, pp. 317–330. 10.1016/B978-0-12-415931-0.00017-3

33. Olson, D.G., Tripathi, S.A., Giannone, R.J., Lo, J., Caiazza, N.C., Hogsett, D.A., Hettich, R.L., Guss, A.M., Dubrovsky, G., Lynd, L.R., 2010. Deletion of the Cel48S cellulase from *Clostridium thermocellum*. Proc. Natl. Acad. Sci. 107, 17727–17732. 10.1073/pnas.1003584107

34. Olson, D. G., 2011. Genetic investigations of the Clostridium thermocellum cellulosome. PhD thesis. Dartmouth College, Hanover, NH.

35. Paye, J.M.D., Guseva, A., Hammer, S.K., Gjersing, E., Davis, M.F., Davison, B.H., Olstad, J., Donohoe, B.S., Nguyen, T.Y., Wyman, C.E., Pattathil, S., Hahn, M.G., Lynd, L.R., 2016. Biological lignocellulose solubilization: comparative evaluation of biocatalysts and enhancement via cotreatment. Biotechnol. Biofuels 9, 8. 10.1186/s13068-015-0412-y

36. Peteranderl, R., Shotts, E.B., Wiegel, J., 1990. Stability of antibiotics under growth conditions for thermophilic anaerobes. Appl. Environ. Microbiol. 56, 1981–1983. 10.1128/aem.56.6.1981-1983.1990

37. Riley, L.A., Ji, L., Schmitz, R.J., Westpheling, J., Guss, A.M., 2019. Rational development of transformation in *Clostridium thermocellum* ATCC 27405 via complete methylome analysis and evasion of native restriction–modification systems. J. Ind. Microbiol. Biotechnol. 46, 1435–1443. 10.1007/s10295-019-02218-x

38. Sanford, P.A., Woolston, B.M., 2022. Expanding the genetic engineering toolbox for the metabolically flexible acetogen *Eubacterium limosum*. J. Ind. Microbiol. Biotechnol. 49, kuac019. 10.1093/jimb/kuac019

39. Schwarz, S., Kehrenberg, C., Doublet, B., Cloeckaert, A., 2004. Molecular basis of bacterial resistance to chloramphenicol and florfenicol. FEMS Microbiol. Rev. 28, 519–542. 10.1016/j.femsre.2004.04.001

40. Seo, H., Lee, J.-W., Garcia, S., Trinh, C.T., 2019. Single mutation at a highly conserved region of chloramphenicol acetyltransferase enables isobutyl acetate production directly from cellulose by Clostridium thermocellum at elevated temperatures. Biotechnol. Biofuels 12, 245. 10.1186/s13068-019-1583-8

41. Silvia, S., Donahue, S.A., Killeavy, E.E., Jogl, G., Gregory, S.T., 2021. A Survey of Spontaneous Antibiotic-Resistant Mutants of the Halophilic, Thermophilic Bacterium Rhodothermus marinus. Antibiotics 10, 1384. 10.3390/antibiotics10111384

42. Su, C.-C., Yin, L., Kumar, N., Dai, L., Radhakrishnan, A., Bolla, J.R., Lei, H.-T., Chou, T.-H., Delmar, J.A., Rajashankar, K.R., Zhang, Q., Shin, Y.-K., Yu, E.W., 2017. Structures and transport dynamics of a Campylobacter jejuni multidrug efflux pump. Nat. Commun. 8, 171. 10.1038/s41467-017-00217-z

43. Suzuki, H., Taketani, T., Kobayashi, J., Ohshiro, T., 2018. Antibiotic resistance mutations induced in growing cells of Bacillus-related thermophiles. J. Antibiot. (Tokyo) 71, 382–389. 10.1038/s41429-017-0003-1

44. Taylor, M.P., Van Zyl, L., Tuffin, I.M., Leak, D.J., Cowan, D.A., 2011. Genetic tool development underpins recent advances in thermophilic whole-cell biocatalysts. Microb. Biotechnol. 4, 438–448. 10.1111/j.1751-7915.2010.00246.x

45. Tian, L., Conway, P.M., Cervenka, N.D., Cui, J., Maloney, M., Olson, D.G., Lynd, L.R., 2019. Metabolic engineering of Clostridium thermocellum for n-butanol production from cellulose. Biotechnol. Biofuels 12, 186. 10.1186/s13068-019-1524-6

46. Tripathi, S.A., Olson, D.G., Argyros, D.A., Miller, B.B., Barrett, T.F., Murphy, D.M., McCool, J.D., Warner, A.K., Rajgarhia, V.B., Lynd, L.R., Hogsett, D.A., Caiazza, N.C., 2010. Development of *pyrF-* Based Genetic System for Targeted Gene Deletion in *Clostridium thermocellum* and Creation of a *pta* Mutant. Appl. Environ. Microbiol. 76, 6591–6599. 10.1128/AEM.01484-10

47. Wada, K., Kobayashi, J., Furukawa, M., Doi, K., Ohshiro, T., Suzuki, H., 2016. A thiostrepton resistance gene and its mutants serve as selectable markers in *Geobacillus kaustophilus* HTA426. Biosci. Biotechnol. Biochem. 80, 368–375. 10.1080/09168451.2015.1079478

48. Walker, J.E., Lanahan, A.A., Zheng, T., Toruno, C., Lynd, L.R., Cameron, J.C., Olson, D.G., Eckert, C.A., 2020. Development of both type I–B and type II CRISPR/Cas genome editing systems in the cellulolytic bacterium Clostridium thermocellum. Metab. Eng. Commun. 10, e00116. 10.1016/j.mec.2019.e00116

49. Xu, Q., Resch, M.G., Podkaminer, K., Yang, S., Baker, J.O., Donohoe, B.S., Wilson, C., Klingeman, D.M., Olson, D.G., Decker, S.R., Giannone, R.J., Hettich, R.L., Brown, S.D., Lynd, L.R., Bayer, E.A., Himmel, M.E., Bomble, Y.J., 2016. Dramatic performance of *Clostridium thermocellum* explained by its wide range of cellulase modalities. Sci. Adv. 2, e1501254. 10.1126/sciadv.1501254

50. Xu, S., Battaglia, L., Bao, X., Fan, H., 2013. Chloramphenicol acetyltransferase as a selection marker for chlamydial transformation. BMC Res. Notes 6, 377. 10.1186/1756-0500-6-377

51. You, Y., Hilpert, M., Ward, M.J., 2013. Identification of Tet45, a tetracycline efflux pump, from a poultry-litter-exposed soil isolate and persistence of tet(45) in the soil. J. Antimicrob. Chemother. 68, 1962–1969. 10.1093/jac/dkt127

52. Zeldes, B.M., Keller, M.W., Loder, A.J., Straub, C.T., Adams, M.W.W., Kelly, R.M., 2015. Extremely thermophilic microorganisms as metabolic engineering platforms for production of fuels and industrial chemicals. Front. Microbiol. 6. 10.3389/fmicb.2015.01209

53. Zhou, J., Wu, K., Rao, C.V., 2016. Evolutionary engineering of *Geobacillus thermoglucosidasius* for improved ethanol production. Biotechnol. Bioeng. 113, 2156–2167. 10.1002/bit.25983

